# Accurate bacterial outbreak tracing with Oxford Nanopore sequencing and reduction of methylation-induced errors

**DOI:** 10.1101/2023.09.15.556300

**Authors:** Mara Lohde, Gabriel E. Wagner, Johanna Dabernig-Heinz, Adrian Viehweger, Sascha D. Braun, Stefan Monecke, Celia Diezel, Claudia Stein, Mike Marquet, Ralf Ehricht, Mathias W. Pletz, Christian Brandt

## Abstract

Our study investigated the effectiveness of Oxford Nanopore Technologies for accurate outbreak tracing by resequencing 33 isolates of a three-year-long *Klebsiella pneumoniae* outbreak with Illumina short read sequencing data as the point of reference.

We detected considerable base errors through cgMLST and phylogenetic analysis of genomes sequenced with Oxford Nanopore Technologies, leading to the false exclusion of some outbreak-related strains from the outbreak cluster. Nearby methylation sites cause these errors and can also be found in other species besides *K. pneumoniae*. Based on this data, we explored PCR-based sequencing and a masking strategy, which both successfully addressed these inaccuracies and ensured accurate outbreak tracing. We offer our masking strategy as a bioinformatic workflow (MPOA is freely available on GitHub under the GNUv3 license: github.com/replikation/MPOA) to identify and mask problematic genome positions in a reference-free manner.

Our research highlights limitations in using Oxford Nanopore Technologies for sequencing prokaryotic organisms, especially for investing outbreaks. For time-critical projects that cannot wait for further technological developments by Oxford Nanopore Technologies, our study recommends either PCR-based sequencing or using our provided bioinformatic workflow. We would advise that read mapping-based quality control of genomes should be provided when publishing results.

## Introduction

Whole genome sequencing is essential for analyzing outbreaks, pandemics, or phylogenetic relationships (Wyres et al. 2020; Chewapreecha et al. 2017). The recent SARS-CoV-2 pandemic has thus led to a leap in the integration and expansion of sequencing capacities in many laboratories and hospitals, predominantly using Illumina for short-read sequencing or Oxford Nanopore Technologies (ONT) for long-read sequencing (approx. 78% and 18%, respectively) (Brandt et al. 2021). Beyond viral pandemic tracking, bacterial pathogen outbreaks, particularly those linked to antibiotic resistance, continue to impose a significant global public health burden (Murray et al. 2022). Gram-negative bacteria, in particular, rapidly acquire antibiotic resistance via horizontal gene transfer from other species (Moura de Sousa et al. 2023; Hadjadj et al. 2022; Lerminiaux and Cameron 2019). This mechanism complicates tracking outbreaks or identifying their origin, as a single specific plasmid or mobile element can be responsible for a persistent outbreak or multiple outbreaks across unrelated species (Hadjadj et al. 2022; Abe et al. 2021; Sivertsen et al. 2014; Pletz et al. 2018). Additionally, in gram-negative bacteria, DNA methylation plays a crucial role in epigenetic regulation, which impacts gene expression, genome modification, virulence, mismatch repair, and other physiological activities (Gao et al. 2023; Wang et al. 2023).

Effectively tracking these complex molecular mechanisms requires careful strategic monitoring and sequencing-based investigation. Consequently, the accuracy and continuity of the genome data are paramount. Illumina, a short-read sequencing method with an error rate of less than 0.8% in raw data, is frequently used as its complementary genome reconstruction precision exceeds 99.997% (Wang et al. 2021). However, repetitive elements, such as transposons, present a substantial challenge for short reads when reconstructing closed bacterial genomes and their accompanying plasmids. Long-read sequencing technologies like Pacific Bioscience (PacBio) and Oxford Nanopore Technologies can resolve such elements, e.g. plasmids, as they achieve longer read lengths averaging around 10-20 kb and even up to 3.85 Mb in the case of ONT (Dohm et al. 2020; Eid et al. 2009; Tyson et al. 2018; Grohme et al. 2018).

Real-time sequencing allows data collection and analysis, while sequencing positions Oxford Nanopore Technologies as an appealing choice for hospital surveillance and outbreak control (Spott et al. 2022). Owing to their recently launched flow cells (R10.4.1) and chemistry (SQK-NBD114.24), they have achieved raw read accuracy that now exceeds 99.1% (Ni et al. 2023). Several studies have shared their findings and reported accuracy levels similar to those from short-read data (Wagner et al. 2023; Sanderson et al. 2023). However, significant discrepancies between Illumina and ONT genomes were also observed for some organisms (Linde et al. 2023).

These contradictions can lead to inaccurate conclusions, like excluding outbreak-associated samples when investigating outbreaks. In addition, genomes are usually stored in open public databases such as NCBI or ENA, which can lead to error propagation and potentially significantly affect patients’ welfare. Therefore, we used ONT to reevaluate a well-documented, three-year-long outbreak initially analyzed with Illumina data to address these contradictory statements, focusing on the errors in ONT sequencing data (Viehweger et al. 2021). *K. pneumoniae* is an ideal microorganism for this topic, as it is a common pathogen linked to hospital-wide outbreaks carrying plasmids with multidrug resistance genes (Brandt et al. 2019). When using ONT-only data, we identified a few critical issues leading to erroneous basecalls for *K. pneumoniae*. We noticed similar problems and clear patterns in other organisms, which need to be considered during outbreak identification, even though we could resolve them.

## Results

### Erroneous basecalls occur in some strains but not others and vary by basecaller and sequencing kits

We resequenced the genomes of 33 randomly distributed *K. pneumoniae* samples isolated from 31 patients (from a total of 114 outbreak-related isolates) using R10.4 and R10.4.1 flow cells, along with the corresponding library preparation kits (henceforth “Kit 12”: SQK-NBD112.24 (early access) and “Kit 14”: SQK-NBD114.24 (successor)). Our objective was to investigate whether previously reported conflicting statements could be replicated (Wagner et al. 2023; Sanderson et al. 2023; Linde et al. 2023). We used core genome multilocus sequence typing (cgMLST) to compare ONT and Illumina-sequenced genomes. The comparison of the 33 samples revealed 11 outliers in the ONT data, showing high allelic deviations (up to 46) to their short-read counterparts while not matching the outbreak cluster (Supplementary Figure 1 and 2). While the remaining samples closely match the outbreak cluster, the outliers highlight inconsistencies within the ONT data, as reported in the literature. To assess whether either the basecaller or their models might be responsible, we re-basecalled and compared an outlier sample (UR2602) in detail to three samples, where we assume error-free genomes based on cgMLST (Figure 1; see method section “Basecalling and Assembly” for further details) and pairwise SNP calling (Supplementary Figure 3) with Illumina genomes.

**Figure 1:**
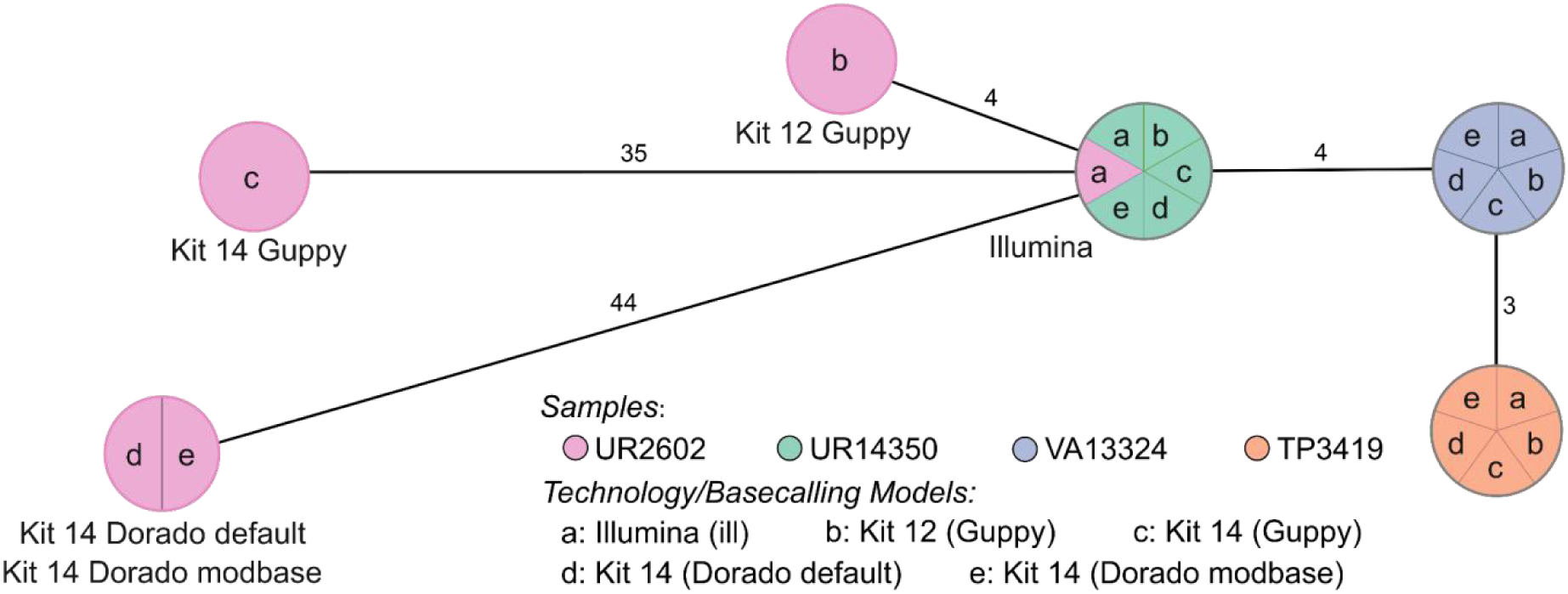
cgMLST typing reveals allelic differences between genomes utilizing different basecaller and models, and sequencing kits. The minimum spanning tree pictures four *K. pneumoniae* samples based on 2,358 genes for pairwise comparison of allelic variations. Missing values were ignored. Nodes (samples) are connected by lines depicting the distance by numbers of allelic differences. Loci are considered different if one or more bases change between the samples. Loci without allelic differences are described as being the same. Samples with allelic differences ≤15 are considered as part of the cluster. All isolates were prepared with Kit 14 and Kit 12 and basecalled with each respective Guppy “super accurate” basecalling model (see methods “Basecalling and Assembly”). We basecalled all Kit 14 - prepared samples with Dorado using the default and a modification-aware model (see methods “Basecalling and Assembly”).

Based on 2,358 loci for the cgMLST, no allelic differences, regardless of kit, basecaller, or basecaller model, were identified for the isolates UR14350, VA13324, and TP3419. In contrast, the outlier sample (UR2602) revealed allelic variations for each kit, basecaller, and sequencing technology. Despite both basecallers using the same raw signal data, the 35 allelic variations in Guppy differ without accordance from the 44 allelic variations in Dorado.

By cgMLST, the outlier sample prepared with Kit 14 would be falsely assigned as not part of the outbreak due to its 35 or 44 allelic differences, even though the sample exhibits low allelic differences according to the short-read data (gold standard). Conversely, when prepared with early access Kit 12, the same isolate would be correctly considered as only four different loci could be observed (adhering to the recommended allelic difference cut-off of ≤15) (Miro et al. 2020). Since the basecalling models disagreed on the allelic differences, we suspected more issues within the raw data (reads and raw signals) and conducted a comprehensive analysis of all possible affected positions.

### Ambiguities in purine or pyrimidine discrimination for a subset of genome positions can cause erroneous basecalls

The first visual inspection of mapped reads to the assembly revealed read ambiguity on certain positions indicated by varying base ratios (ambiguous positions). For further characterization of these ambiguous positions, we examined our data on the sequence, nucleotide, and raw signal level (Figure 2). For each ambiguous position on the chromosomal DNA for 33 *K. pneumoniae* outbreak samples, we determined the ratio between the two bases by counting their occurrences for both strand orientations within the read data at that position (Figure 2 A). Searching for characteristic “indicator” sequence motifs, we explored the surrounding base for each detected ambiguous position and plotted the observed pattern as a sequence logo (Figure 2 B and Supplementary Table 1). Additionally, we compared the methylated and unmethylated raw signals around these ambiguous positions (Figure 2 C).

**Figure 2:**
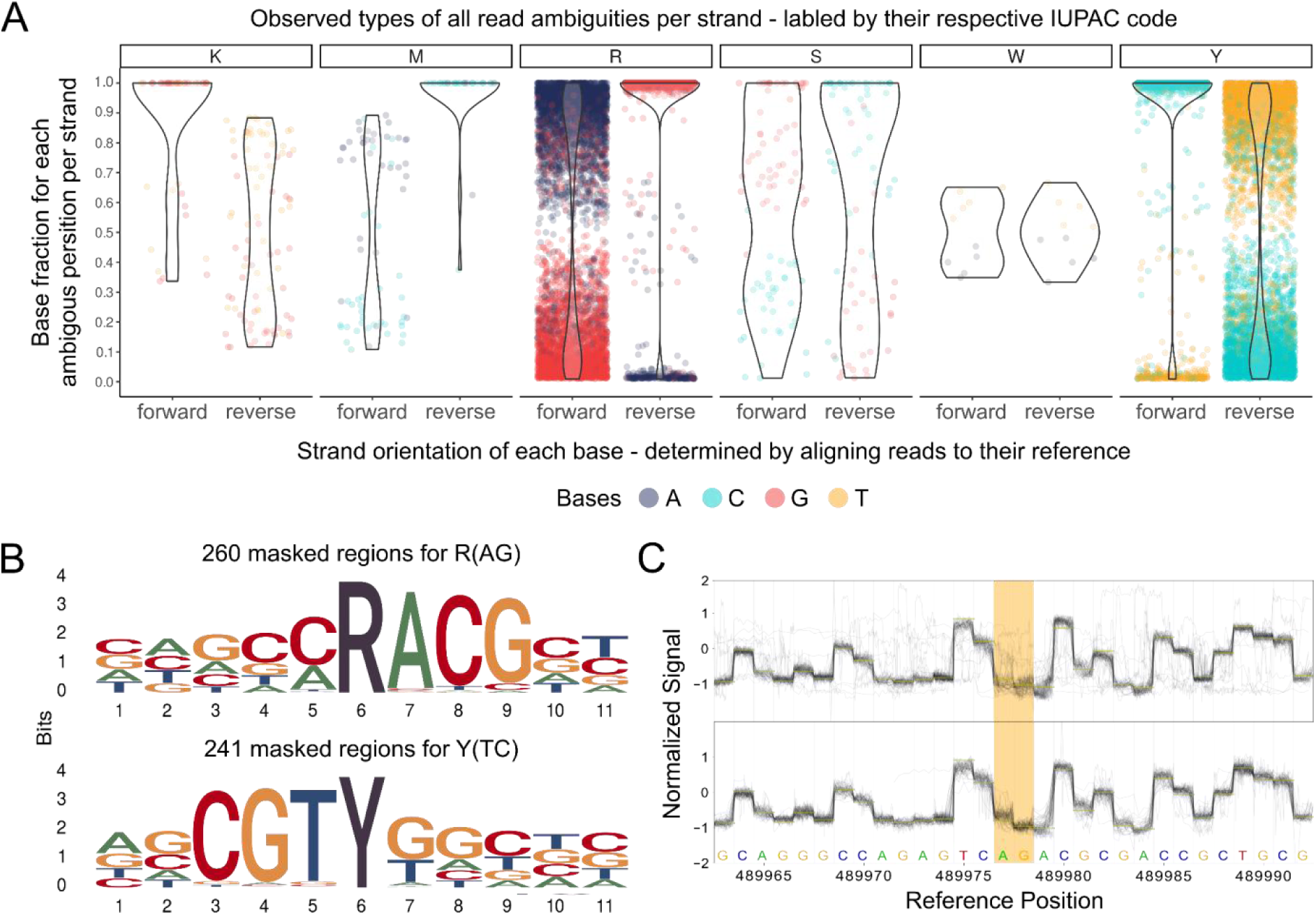
Systematic examination of ambiguous positions for frequency by strand orientation (A), conserved sequences (B), and raw Nanopore signal (C). **A:** Violin chart showing the ratio between two bases within the mapped read data separated by strand orientation for 6,556 ambiguous positions in 33 *K. pneumoniae* samples. Every ambiguous position is divided by which two bases appear and labeled by their respective degenerative base (IUPAC nucleotide code). For example, “R” stands for a combination where either A or G is found at that position. Each dot represents a base occurrence within the respective base combination at the ambiguous position. **B:** Sequence logo of observed sequence pattern around the ambiguous bases R and Y on the chromosomal contig of *K. pneumoniae* for one sample. **C:** Raw signal level (FAST5/POD5) of ambiguous positions (yellow) for Kit 14 (above) with methylated bases and SQK-RPB114.24 without modifications (below). Less clear signals are observable in ambiguous positions (yellow) for Kit 14. Signal plots were generated with remora (v.2.1.3; github.com/nanoporetech/remora).

As highlighted in Figure 2 A, in some positions, the basecaller can not determine between either of two bases, expressed by specific base ratios varying per position and strand orientation, resulting in erroneous assemblies. We could not observe ambiguous positions containing more than two different bases. For clarity, we assign the IPUAC nucleotide code for degenerate bases (R, Y, M, K, S, W) to each ambiguous position varying between two bases. Accordingly, we will refer, e.g. to “R” when the positions contain A or G in the read data.

Out of our analysis of 33 *K. pneumoniae* outbreak isolates, we discovered 6,556 positions that exhibit ambiguity (Figure 2A). The ambiguity mainly resolved around 3,311 positions for R and 3,111 for Y. In 5,442 of 6,455 R and Y positions (84.31%), the basecalled reads lean towards Cytosine or Guanine (C or G). We detected other ambiguous positions in K (44), M (34), S (51), and W (5), but with comparable lower occurences. It is essential to acknowledge that not all identified ambiguous positions result in errors in the assembled genome, which explains the varying error profile of the same sample (Figure 1). Errors in these ambiguous positions mainly arise when deciding between purine bases (A or G) or pyrimidine bases (T or C). Since strand bias is reported as a substantial factor for false positive variant detection, we considered the strand orientation of the read data (Leija-Salazar et al. 2019). In most cases, we noticed that the correct base is located clearly and more frequently on both strand orientations, while the incorrect base is less prevalent. For instance, the correct base Guanine is found on both strands for positions masked by R and is predominantly found on the reverse strand (Fig 2 A: see R reverse), less frequently on the forward strand, which also contains the false base. We detected similar behavior for other species (Supplementary Figure 6).

In the error-prone genomes of the *K. pneumoniae* outbreak, we detected preserved patterns around the ambiguous positions R and Y (Figure 2B). These sequence motifs are reverse-complement patterns (RACG/CGTY), pointing to a singular issue. Compared to other isolates of *K. pneumoniae*, we also observed additional patterns. These motifs are likely specific to particular strains.

Furthermore, we examined and collected additional ONT sequencing data that used the Kit 14 library preparation (264 isolates across 32 species) to investigate whether the ambiguous positions are *K. pneumoniae* exclusive (Table 1). These samples were collected based solely on Kit 14 library preparations and not on whether they were associated with an outbreak. Compared to all other species samples, we determine the fewest ambiguous positions (0 to 1) in *Bordetella pertussis*. In contrast, all 10 *Enterococcus faecalis* isolates had over 200 ambiguous positions. Over 40% of 264 screened samples have more than 50 ambiguous positions. Across all species, the minimal shared sequence motif was RA/TY. Certain species, such as *Acinetobacter junii*, *Acinetobacter radioresistens*, *Chryseobacterium gleum*, *Enterobacter cloacae*, *Micrococcus luteus,* and *Stenotrophomonas maltophilia*, exhibited a considerable number of ambiguous positions. This suggests that many species may be impacted, but not necessarily all strains.

**Table 1:** Overview of R (A-G) and Y (T-C) base ambiguity for 264 isolates from 32 various species, sequenced with Oxford Nanopore Technologies using Kit 14. Only chromosomal contigs were analyzed, and only “super accurate” basecalling models were used. Chromosomes were coverage-masked by N if below a read depth of 10x. N positions were not considered for the table to avoid overestimating one base ambiguity.

| Species | Total samples | Samples with > 50 ambig. P. | Mean R (A-G) ambiguity per sample (min/max) | Mean Y (T-C) ambiguity per sample (min/max) | Motif type | Reference |
| --- | --- | --- | --- | --- | --- | --- |
| *Achromobacter xylosoxidans* | 1 | 0 (0%) | 6 | 4 | NA | * |
| *Acinetobacter baumannii* | 15 | 5 (33.33%) | 22.53 (0/109) | 19.87 (0/83) | NA | * |
| *Acinetobacter junii* | 1 | 1 (100%) | 279 | 223 | CARATG<br>CATYTG | * |
| *Acinetobacter mesopotamicus* | 1 | 1 (100%) | 53 | 28 | NA | * |
| *Acinetobacter radioresistens* | 1 | 1 (100%) | 175 | 150 | RA<br>TY | * |
| *Acinetobacter soli* | 1 | 0 (0%) | 12 | 11 | NA | * |
| *Bordetella pertussis*<sup>§</sup> | 40 | 0 (0%) | 0.1 (0/1) | 0.2 (0/1) | NA | (Wagner et al. 2023) |
| *Chryseobacterium arthrosphaerae* | 1 | 0 (0%) | 13 | 7 | NA | * |
| *Chryseobacterium gleum* | 1 | 1 (100%) | 218 | 217 | RACGC<br>GCGTY | * |
| *Citrobacter freundii* | 3 | 3 (100%) | 145.67 (49/329) | 137.67 (39/319) | CRATGTC<br>GACATYG | * |
| *Citrobacter portucalensis* | 2 | 2 (100%) | 29 (26/32) | 29 (28/30) | RA<br>TY | * |
| *Enterobacter cloacae* | 1 | 1 (100%) | 153 | 162 | NA | * |
| *Enterobacter hormaechei* | 5 | 1 (20%) | 15.60 (4/45) | 15.60 (6/42) | NA | * |
| *Enterococcus faecalis* | 10 | 10 (100%) | 250.40 (223/275) | 249.00 (210/270) | TRAG<br>CTYA | + |
| *Enterococcus faecium* | 19 | 2 (10.53%) | 15.74 (0/27) | 14.63 (0/28) | RACC<br>GGTY | # |
| *Escherichia coli* | 8 | 3 (37.50%) | 31.38 (2/118) | 29.31 (4/111) | NA | * |
| *Escherichia flexneri* | 10 | 9 (90%) | 49.10 (19/88) | 44.70 (18/88) | RAT<br>ATY | * |
| *Klebsiella aerogenes* | 1 | 0 (0%) | 22 | 17 | NA | * |
| *Klebsiella michiganensis* | 1 | 1 (100%) | 56 | 42 | NA | * |
| *Klebsiella pneumoniae* | 70 | 38 (54.29%) | 97.04 (3/835) | 92.47 (3/847) | RACG<br>CGTY | (Viehwe<br>r et al.<br>2021)<br>* # |
| *Klebsiella oxytoca* | 1 | 0 (0%) | 5 | 9 | NA | * |
| *Listeria monocytogenes* | 17 | 3 (17.65%) | 37.94 (0/514) | 40.82 (0/557) | NA | # |
| *Micrococcus luteus* | 1 | 1 (100%) | 172 | 220 | CRAC<br>GTYG | * |
| *Proteus mirabilis* | 1 | 1 (100%) | 45 | 43 | CRAC<br>GTYG | * |
| *Pseudomonas aeruginosa* | 19 | 6 (33.33%) | 389.17 (0/2251) | 387.56 (0/2290) | AARACC<br>GGTYTT | * |
| *Pseudomonas asiatica* | 2 | 1 (50%) | 145 (0/290) | 144.50 (0/289) | CCRA<br>TYGG | * |
| *Pseudomonas stutzeri* | 1 | 0 (0%) | 14 | 16 | NA | * |
| *Salmonella enterica* | 2 | 1 (50%) | 41.5 (7/76) | 42.5 (7/78) | NA | * |
| *Stenotrophomonas maltophilia* | 1 | 1 (100%) | 229 | 171 | TACRAC<br>GTYGTA | * |
| *Serratia marcescens* | 4 | 4 (100%) | 107.75 (68/178) | 99.5 (67/171) | CCRA<br>TYGG | * |
| *Shewanella algae* | 2 | 2 (100%) | 71.5 (37/106) | 50 (32/68) | NA | * |
| *Staphylococcus aureus* | 20 | 8 (40%) | 23.35 (0/99) | 23.35 (0/97) | RACC<br>GGTY | # |

\* Sequenced strains received from Leibniz-Institute of Photonic Technology, Optisch-Molekulare Diagnostik und Systemtechnologie

As methylated bases are probably liable for ambiguous positions, we compared sequencing data with methylations (Kit 14) (Figure 2 C above) and without (SQK-RPB114.24) (Figure 2 C below) on the raw signal level from FAST5/POD5 files before basecalling occurs. We choose the raw signal level for investigation prior to any bioinformatic approaches (e.g., basecalling, assembly, polishing) to avoid the potential introduction of other biases or errors. For native sequencing, less clear signals at these positions are observable, which might cause these ambiguous basecalls. These noisy signals could explain the frequencies of bases we detected in the reads (Figure 2A) and, thus, the basecaller’s difficulty in deciding on a specific base for that position.

We found no coherent methylation motifs in the literature that would fit the observed pattern. Nevertheless, it has been reported that methylated bases can affect the raw signal in the surrounding region (Tourancheau et al. 2021). Thus, we can not determine whether multiple methylation motifs are the cause or if an unknown motif is present. Accordingly, to these findings, we evaluated whether PCR-based sequencing or a bioinformatic masking strategy for ambiguous positions can reliably remove these methylation-based errors for outbreak analysis.

### Strategies to mitigate methylation-induced basecalling errors

To solve methylation-induced basecalling errors in ambiguous base positions, we evaluated two strategies: 1) We resequenced 10 *K. pneumoniae* outbreak samples using the Nanopore Rapid PCR Barcoding Kit (SQK-RPB114.24) to remove methylated bases prior to sequencing and analyzed the genomes using cgMLST and phylogenetic analysis (Figure 3 A and B). 2) We masked ambiguous positions for Kit 14 prepared genomes with our bioinformatic workflow (see method section “Workflow for detecting and masking of ambiguous positions”). It is important to mention that these masked assemblies cannot be used for cgMLST because allelic differences cannot be accurately determined for genes with masked bases. Therefore, the masked genomes were only used for phylogenetic analysis (Figure 3 B). Furthermore, it should be noted that cgMLST only considers coding sequences, whereas the phylogenetic analysis assesses the entire genome that is represented in all samples, which gives a higher resolution for base differences.

**Figure 3:**
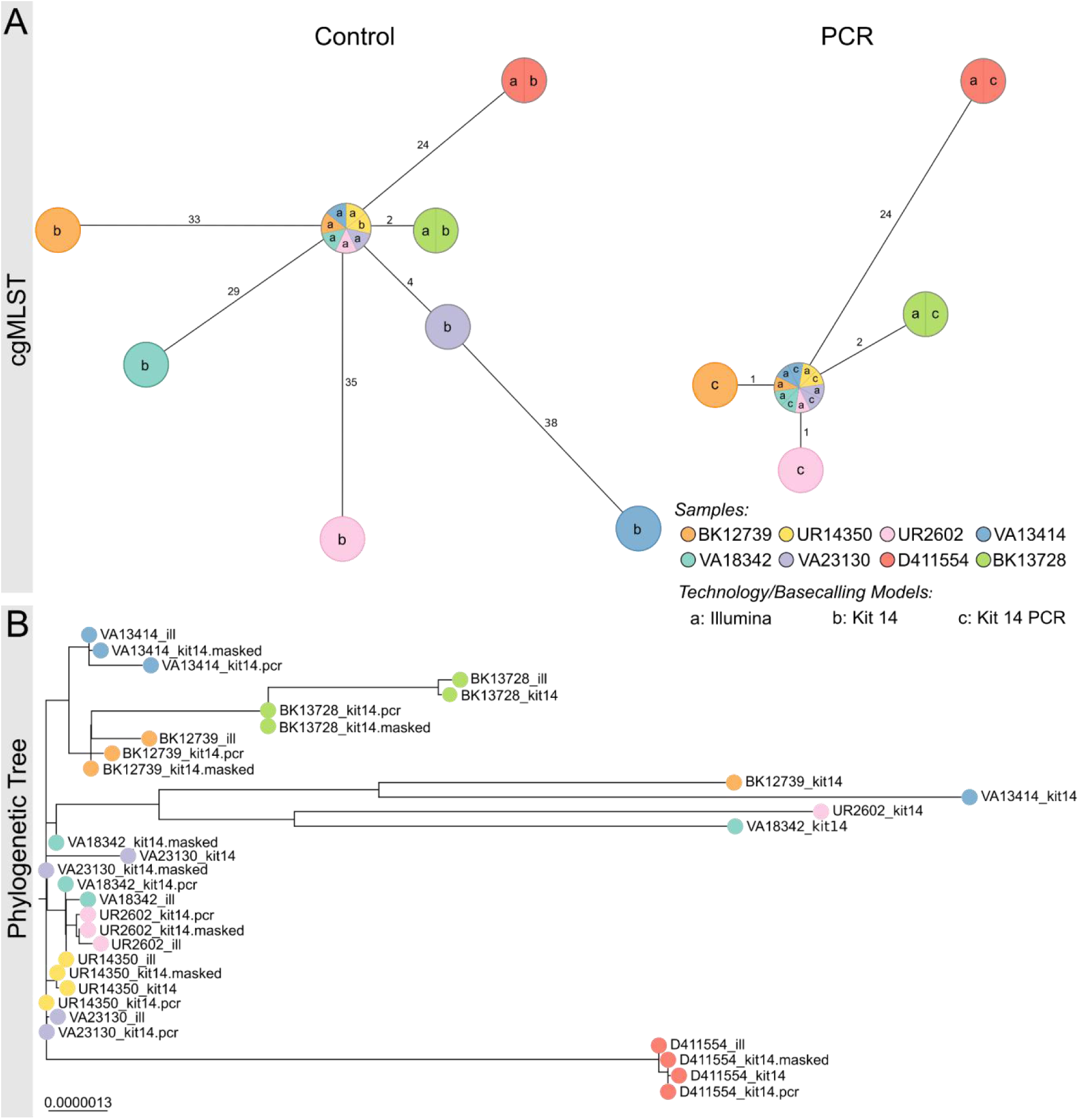
PCR-based sequencing or masking of ambiguous positions reduces allelic or phylogenetic distances. **A:** Minimum spanning trees (pairwise ignore missing values) of each of eight *K. pneumoniae* outbreak samples based on 2,358 genes to compare allelic differences between Illumina and Nanopore SQK-NBD114.24 genomes (kit14; left) and Illumina and Nanopore SQK-RPB114.24 genomes (PCR; right). Nodes (samples) are connected by lines depicting the distance by numbers of allelic differences. Loci are considered different whether one or more bases change between the samples. Loci without allelic differences are described as being the same. Samples with allelic differences ≤15 are considered as part of the cluster. **B:** Phylogenetic tree based on core genome SNP alignment between eight *K. pneumoniae* outbreak samples (colored nodes), prepared with Illumina (ill), Nanopore SQK-NBD114.24 (kit14) and SQK-RPB114.24 (pcr) compared to the masked Kit 14 assemblies (masked).

When comparing the Native Barcoding with the PCR-based Kit, ambiguous positions were significantly reduced from 2,316 to just 14 for R and Y across the 10 resequenced *K. pneumoniae* samples (Supplementary Table 2). Both the minimum spanning trees and the phylogenetic tree also show this significant improvement in genome quality for ONT (Figure 3). According to the cgMLST, the outlier samples UR2602 and BK12739 now closely match the Illumina genome, down to only one allele difference from 35/33 (Figure 3 A). When comparing phylogenetic distances within the phylogenetic tree, an increased convergence with the Ilumina genomes, particularly for the outlier samples, was observed, too (Figure 3 B). Additionally, masked and PCR-based assemblies have almost no phylogenetic divergence.

Further, we analyzed the phylogenetic tree containing native Kit 14, masked native Kit 14, and Illumina genomes for all 33 *K. pneumoniae* samples (Supplementary Figure 4). These include 11 Kit 14 outliers (average of 492 ambiguous positions) and 22 Kit 14 genomes with an average of less than 52 ambiguous positions. We observed two types of phylogenic distances between ONT Kit 14 and Illumina: The expected considerable distances between the outlier and Illumina genomes are due to ambiguity and, in some cases, a phylogenetic distance for which ambiguity is not the causation.

By masking ambiguous bases, we observed that 8 of 11 outlier genomes now closely align with their respective Illumina genome. The remaining three outlier samples changed their tree positions after masking, now closely aligning with other Illumina genomes but still diverged from their corresponding Illumina genome due to other non-ambiguity-related differences. For the other 22 masked Kit 14 genomes with less ambiguity than the outliers, we did not observe any substantial changes in their tree positions, as fewer positions were masked.

In summary, 22 out of 33 masked ONT genomes align with their respective Illumina genomes, and the remaining 11 do not. In these cases, the remaining distances do not result from ambiguous positions within the ONT assemblies. For instance, the Illumina and ONT genomes of TP3870 matched perfectly in the minimum spanning tree, but they exhibited some distance from each other in the phylogenetic tree (Supplementary Figure 4). We identified reconstruction issues in these short-read assemblies, primarily manifesting in non-coding regions (Supplementary Figure 5). Since cgMLST only compares coding sequences, these errors do not affect the result analysis. Therefore, we recommend using only one technology when performing whole genome comparison for outbreak analysis.

## Discussion

Over the past few years, Oxford Nanopore Technologies has been effectively used to monitor and track the SARS-CoV-2 pandemic and its viral lineages. Despite this, contradictory reports have emerged regarding the consistency of ONT-sequenced bacterial genomes compared to Illumina-based. Our research examined whether Oxford Nanopore Technology can be used to analyze bacterial outbreaks accurately.

For our investigation, we resequenced a well-documented 3-year *K. pneumoniae* outbreak using the Nanopore Native Barcoding Kit 14 for library preparation. Our analyses demonstrated that the raw signals were impacted by methylated bases, creating ambiguous positions through basecalling and leading to erroneous exclusions of certain outbreak-associated strains. However, not all isolates are affected by these ambiguities, and none or minimal allelic differences are shown in the corresponding short-read data. One should note that some errors in non-coding areas for Illumina assemblies were observed, which can lead to higher distances within a phylogenetic tree but with little to no effect in cgMLST. Despite focusing on *K. pneumoniae* initially, other prokaryotic organisms are also impacted. Crucially, we also detected ambiguous positions using the predecessor sequencing Kit SQK-LSK109 (Supplementary Table 3). Consequently, when using ONT-based sequence data from open public databases or when analyzing outbreaks, test for read ambiguity by using e.g. the provided MPOA workflow before further analysis.

Based on our in-depth investigation, we recommend using the Nanopore Rapid PCR Barcoding Kit for sequencing to eliminate these read ambiguities in the genome assemblies. However, this method decreases the read length to roughly 3,500 bp, posing difficulties in achieving closed plasmids and genomes, similar to other short-read approaches but to a way lesser extent. A higher sequencing depth might also be necessary to control for polymerase errors. For samples already sequenced without any involvement of PCR, we propose using the provided MPOA workflow to assess the quality of each genome. This workflow offers information about the frequency and strand orientation of reads in ambiguous positions and masks them in the assembly by the IUPAC nucleotide code without needing another reference. These masked assemblies can be used for constructing phylogenetic trees for outbreak tracking. However, cgMLST cannot be performed as masked or degenerated bases create false allelic differences across the whole minimal spanning tree.

Given the notable strides made in direct methylation calling techniques, Oxford Nanopore Technology might overcome the issues with ambiguous positions. If available in high enough quantities, duplex reads (connecting and sequencing both strands) might provide better raw signal data for accurate basecalling. The recently introduced research model reduces the ambiguous positions in direct comparison, but still a few remain (Supplementary Table 4; Researchmodel “res_dna_r10.4.1_e8.2_400bps_sup” available on github.com/nanoporetech/rerio, not yet implemented in MinKnow version 23.07.12, as of 30.11.2023). Further advances that reduce methylation-induced errors for certain specific motifs are being developed, as shown for *Listeria monocytogenes* and *Escherichia coli* (Chiou et al. 2023; Hallgren et al. 2021). Nevertheless, we strongly recommend constantly testing and evaluating reconstructed prokaryotic genomes to avoid erroneous conclusions based on these ambiguous positions introduced by unknown and not yet considered methylation motifs.

## Methods

### Isolates and Genomic Data

Oxford Nanopore Technologies sequencing data from three institutes have been collected and analyzed. The sequencing data includes 264 isolates from 32 species, provided by the Leibniz-Institute of Photonic Technology Jena, Medical University of Graz, and University Hospital Jena. Additionally, a set of 80 samples containing *K. pneumoniae*, *E. faeces*, *L. monocytogenes*, *and Staphylococcus aureus* from a ring trail were used for analysis. University Hospital Leipzig provided 33 Carbapenem-resistant *K. pneumoniae* outbreak isolates from sequence type 258 and the associated Illumina sequencing data (Supplementary Table 5). We ensured these samples were scattered across the whole phylogenetic tree that Viehweger et al. provided in their manuscript.

### Genomic DNA Isolation

Isolates from 10% glycerin cryo culture streaked out on Columbia Agar with 5% Sheep Blood (Becton Dickinson). After overnight incubation, a single colony was selected and cultured overnight in MH-Broth. Genomic DNA was isolated via ZymoBIOMICS DNA Microprep Kit (D4301 and D4305; ZymoResearch) with modifications to enhance the output yield. Qubit dsDNA BR Assay-Kit (Thermo Fisher Scientific) was employed to quantify DNA concentrations from each sample. This kit uses fluorescent dyes to measure double-stranded DNA to ensure reliable results.

### Whole Genome Sequencing

To prepare the library for sequencing using Oxford Nanopore Technologies’ GridION system, we used the Native Barcoding Kit 24 V12 (SQK-NBD112.24, Oxford Nanopore Technologies) and Native Barcoding Kit 24 V14 (SQK-NBD114.24, Oxford Nanopore Technologies) with R10.4 and R10.4.1 flow cells, respectively. Both sequencing protocols were optimized regarding prolonged incubation times. Additionally, one library was prepared with Rapid PCR Barcoding Kit 24 (SQK-RPB114.24, Oxford Nanopore Technologies) for sequencing on an R10.4.1 flow cell. Sequencing of libraries prepared with SQK-NBD112.24 and SQK-NBD114.24 was conducted at 4kHz with 260bps, while SQK-RPB114.24 was conducted at 5kHz with 400bps. The DNA fragments minimum length for all sequencing runs was set to 200bp in MinKNOW (v22.12.5) software.

### Basecalling and Assembly

Basecalling and barcode demultiplexing of long-read sequencing data was performed on the GridiION (Oxford Nanopore Technologies) deploying Guppy (v6.4.6) using super accurate models associated with the different used sequencing kits (“dna_r10.4_e8.1_sup.cfg”, “dna_r10.4.1_e8.2_260bps_sup.cfg”, “dna_r10.4.1_e8.2_5khz_400bps_sup.cfg”). For further analysis, Dorado (v0.3.0) was used (“dna_r10.4.1_e8.2_260bps_sup.cfg” and “dna_r10.4.1_e8.2_260bps_modbases_5mc_cg_sup.cfg”).

Reads were filtered by excluding them below 1,000bp with Filtlong (v.0.2.1) (https://github.com/rrwick/Filtlong). *De novo* assembly was conducted using Flye (--meta --nano-hq) (v2.9) (Kolmogorov et al. 2019). Filtered reads were mapped to the assembly using Minimap2 (-ax map-ont) (v2.18) (Li 2018) and polished afterward by Racon (v1.4.20; github.com/lbcb-sci/racon) followed by medaka_consensus polishing both in default settings (v1.5.0; github.com/nanoporetech/medaka) using the following models: *r104_e81_sup_g5015*, *dna_r10.4.1_e8.2_260bps_sup@v3.5.2*, *r1041_e82_260bps_sup_g632* (Supplementary Code 1). Short reads obtained by Illumina sequencing were assembled using Shovill (v1.1.0, github.com/tseemann/shovill), which includes various genome corrections and polishing approaches (Viehweger et al. 2021).

### Core genome multilocus sequence typing of *K. pneumonia*

For core genome multilocus sequence typing (cgMLST), we used a species-specific public cgMLST scheme for a gene-by-gene comparison on an allelic level (Mellmann et al. 2016; Leopold et al. 2014) according to the ‘*K. pneumoniae sensu lato* cgMLST’ (https://www.cgmlst.org/ncs/schema/2187931/). This comprises a total of 2358 genes (about 40% of the NTUH-K2044 reference genome) (Bialek-Davenet et al. 2014) in flye-assembled and polished long-read and Shovill-assembled short-read assemblies (v1.1.0, github.com/tseemann/shovill). To illustrate the clonal relationships between different isolates, a minimum-spanning tree analysis was performed based on the determined allelic profiles using the Ridom SeqSphere^+^ software version 7 (Ridom GmbH, Muenster, Germany) (Jünemann et al. 2013) with the parameter “pairwise ignore missing values”. We defined a clonal transmission event if the isolates differ ≤15 alleles for *K. pneumoniae* (Miro et al. 2020).

### Phylogenetic tree

The phylogenetic trees visualize the evolutionary relationship among *K. pneumoniae* outbreak strains and are constructed based on core genome SNP alignment using Snippy-core (v4.6.0; github.com/tseemann/snippy). The phylogenetic tree was built using FastTree (Price et al. 2009) and visualized in Microreact (Argimón et al. 2016).

### Workflow for detection and masking of ambiguous positions

We developed a standardized Nextflow (Di Tommaso et al. 2017) workflow for *de novo* quality validation of all species, which is publicly available at github.com/replikation/MPOA, licensed under GNU General Public License v3.0. The workflow only needs the genome file (FASTA) and the associated reads (FASTQ) (Figure 4). The workflow provides reproducible quality control by counting and summarizing ambiguous bases for the user, masking low coverage regions (0-10x depth) with BEDTools (Quinlan and Hall 2010) (v2.31.0), and providing an assembly with these positions masked by the IUPAC nucleotide code for subsequent analysis. The workflow utilizes Docker or singularity container and is compatible with a local run, Slurm, or Google Cloud Compute. Identification and masking of ambiguous positions were conducted using SAMtools consensus (Li et al. 2009) (v1.17) after Minimap2 (default) (v.2.26) (Li 2018) or BWA (v.0.7.17) (alternative) mapping (Li and Durbin 2010) (v.2.26). Different mapping approaches were tested (Supplementary Figures 7 and 8). PlasFlow (Krawczyk et al. 2018) (v1.1.0) extracts chromosome contigs for downstream analysis without plasmid sequences. R was utilized to plot sequence motifs (Figure 2B) with ggseqlogo (Wagih 2017) and a violin chart comparing base frequencies per strand (Figure 2A) with ggplot2 (Wickham 2016).

**Figure 4:**
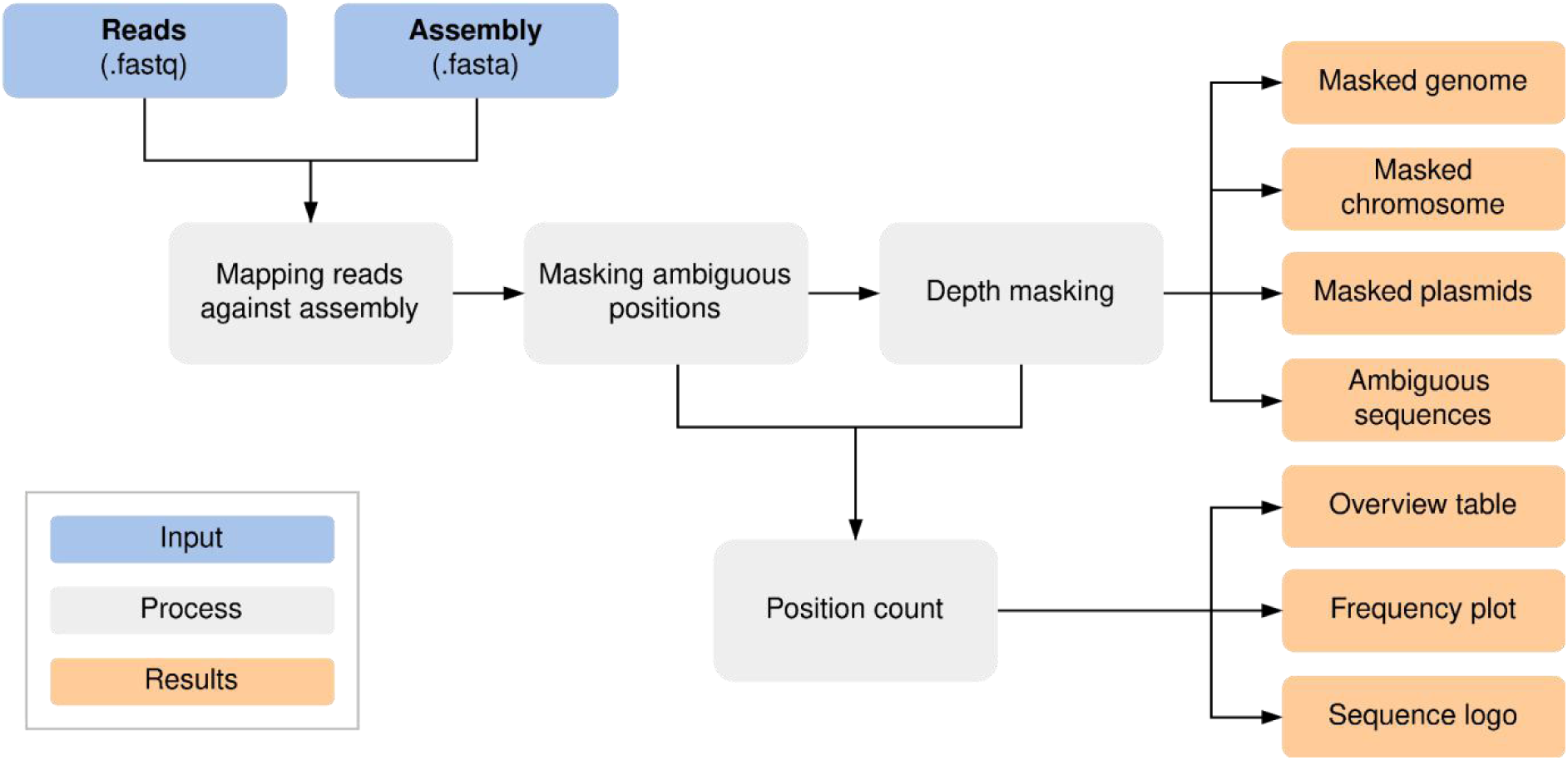
MPOA workflow to mask ambiguous and low coverage (0-10x sequencing depth) positions in genome files. The workflow provides masked assemblies containing all contigs and separate masked chromosomes and plasmid FASTA files. A FASTA file per sample is also generated for each ambiguous position plus surrounding bases for further analysis (github.com/replikation/MPOA; Supplementary Code 2).

## Data access

The Workflow used in this study has been uploaded to GitHub (github.com/replikation/MPOA) and Supplementary_code_2.zip.

The Oxford Nanopore Technologies sequencing data generated in this study have been submitted to the NCBI BioProject database (https://www.ncbi.nlm.nih.gov/bioproject/?term=PRJNA1050168) under NCBI BioProject ID PRJNA1050168.

The Illumina sequencing data used in this study have been submitted to the NCBI BioProject database (https://www.ncbi.nlm.nih.gov/bioproject/?term=PRJNA742413) under NCBI BioProject ID PRJNA123456.

## Acknowledgments

This work received financial support from the Ministry for Economics, Sciences and Digital Society of Thuringia (TMWWDG) under the framework of the Landesprogramm ProDigital (DigLeben-5575/10-9) and the Open Access Publication Fund of the Thueringer Universitaets-und Landesbibliothek Jena.

## Authors’ contributions

Sample collection and preparation: M.L., A.V., C.D; sequencing: M.L.; workflow development and testing: M.L. and C.B.; bioinformatic analysis: M.L. and C.B.; literature research: M.L.; writing first draft: M.L.; reviewing and editing manuscript: M.L., C.B., A.V., C.S, M.M., G.W.L, J.D.H., R.E, S.D.B., S.M., C.D and M.W.P.; supervision: C.B.; project administration: C.B.; cgMLST analysis: C.S.; funding acquisition: C.B.; providing genomes: M.L., S.D.B., S.M., A.V., G.W.L, J.D.H; All authors have read and agreed to the published version of the manuscript.

