## Supplementary data for "Accurate bacterial outbreak tracing with Oxford Nanopore sequencing and reduction of methylation-induced errors"

**Supplementary Tables** 1

**Supplementary Figures** 10

**Supplementary Code** 20

### Supplementary Tables

**Supplementary Table 1:** Sequence logos of observed sequence pattern around the ambiguous bases R and Y on the chromosomal contig for different species based on one sample.

<table>
<tr><th>Species</th><th>Motif type</th></tr>
<tr><td>Acinetobacter junii</td><td>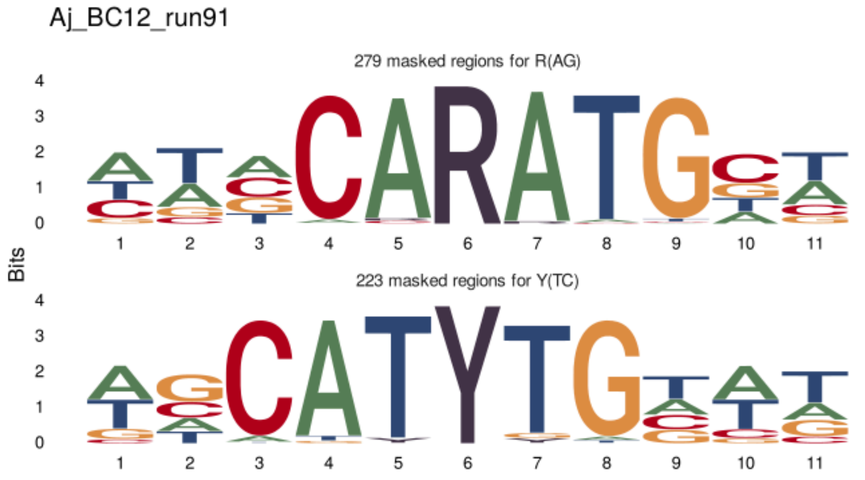
</td></tr>
<tr><td>Acinetobacter radioresistens</td><td>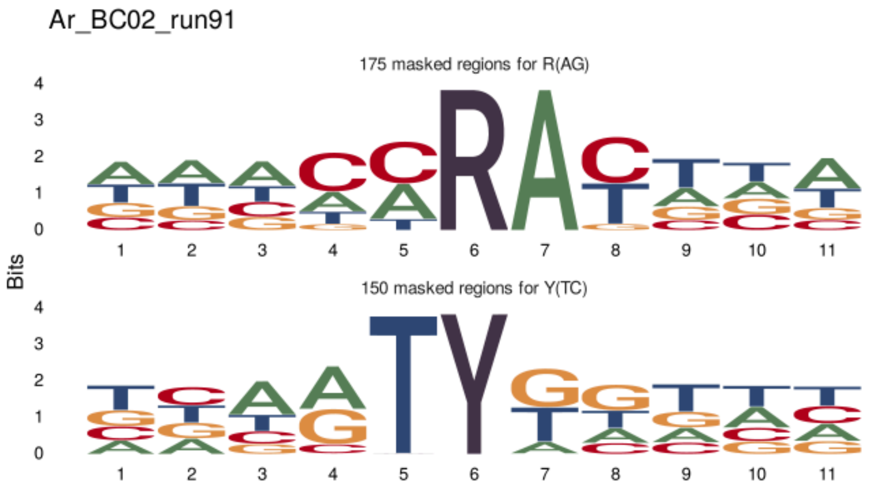
</td></tr>
</table>

*Chryseobacterium gleum*

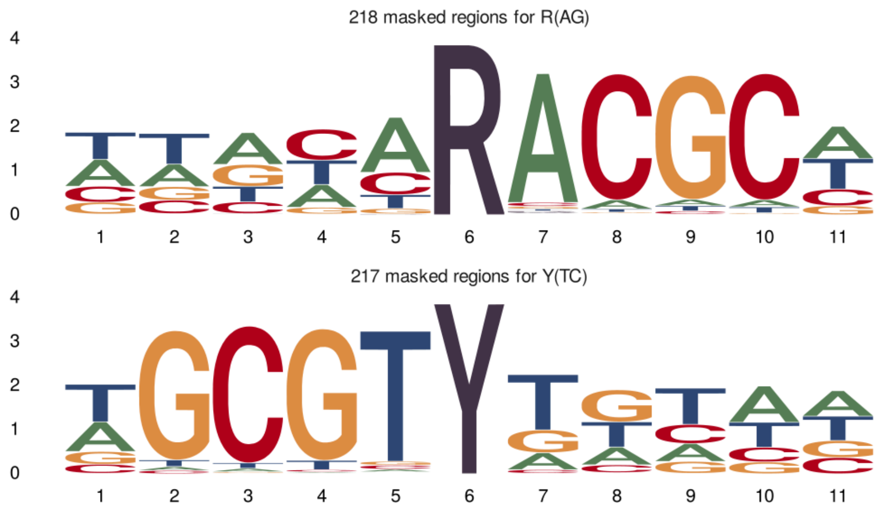


*Citrobacter freundii*

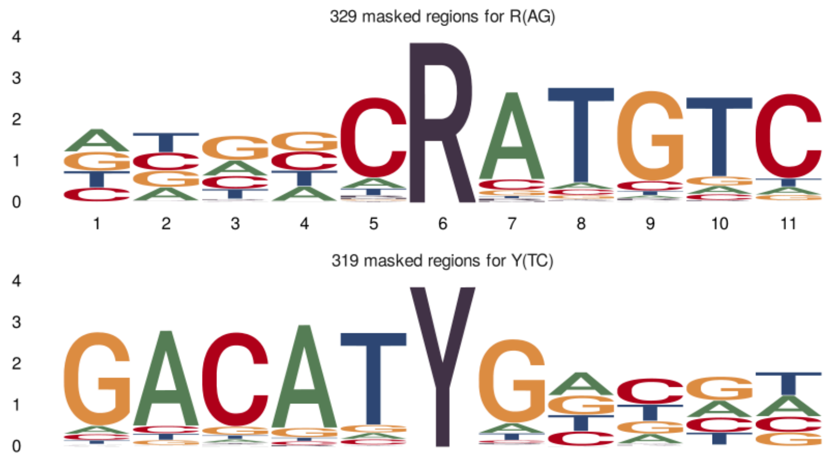


*Enterococcus faecalis*

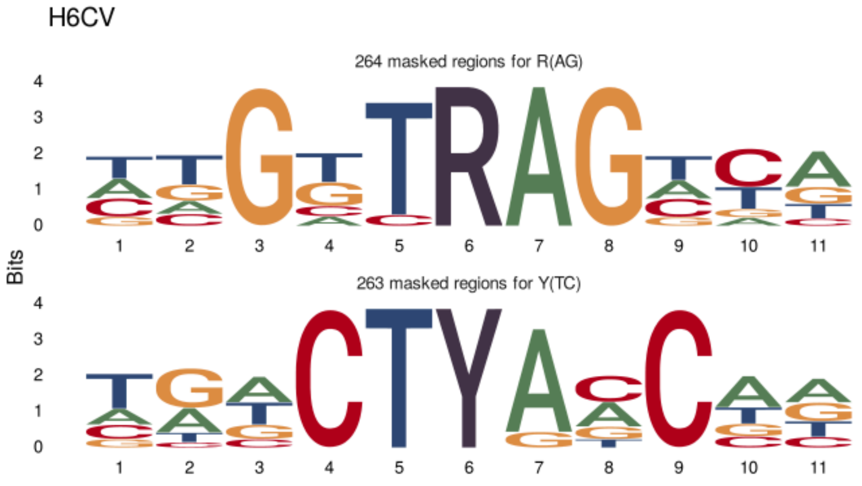

<table>
<tr>
<td>Enterococcus faecium</td>
<td>
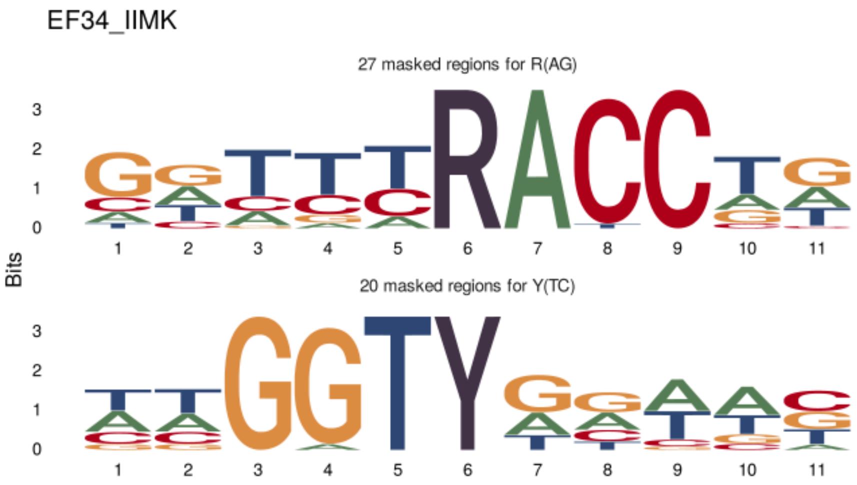

</td>
</tr>
<tr>
<td>Escherichia flexneri</td>
<td>
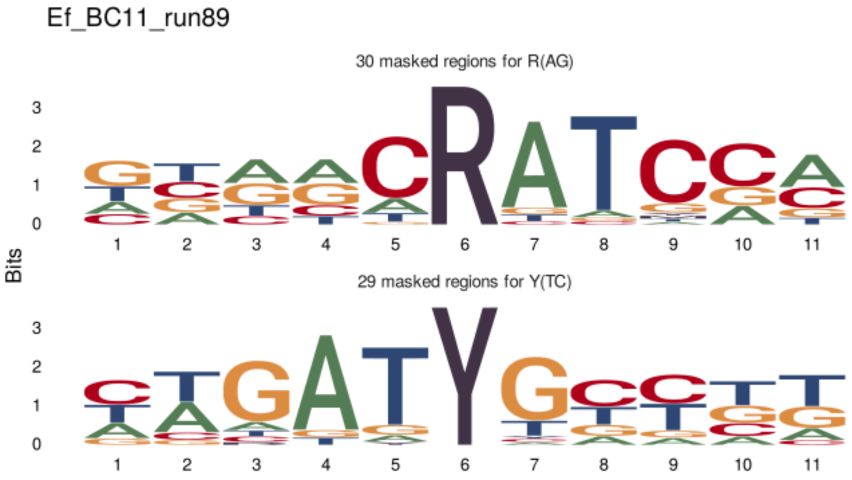

</td>
</tr>
<tr>
<td>Klebsiella pneumoniae</td>
<td>
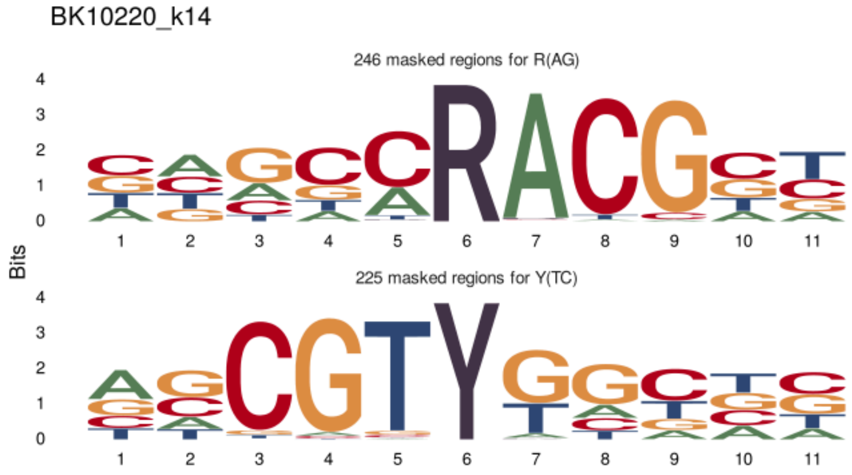

</td>
</tr>
</table>

| | |
|---|---|
| *Serratia marcescens* | 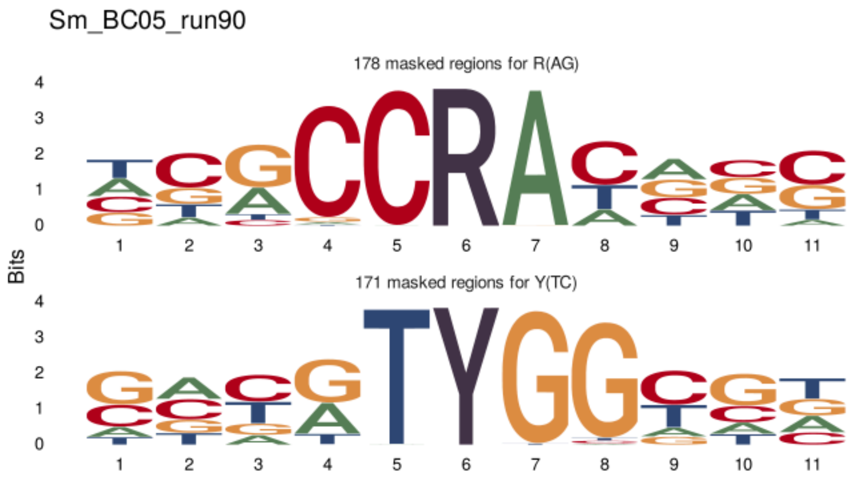 |

**Supplementary Table 2:** Evaluation of errors in reads for different species. Native reads (SQK-NBD114.24) and PCR reads (SQK-RPB114.24) were compared against the same native genome. Only the most problematic observed ambiguous positions, R and Y, within the chromosome, are included in this table. The evaluation was done using MPOA workflow.

| | native reads (SQK-NBD114.24) | | PCR reads (SQK-RPB114.24) | |
|---|---|---|---|---|
| | R (A or G) | Y (T or C) | R (A or G) | Y (T or C) |
| *Klebsiella pneumoniae* (UR2602) | 260 | 241 | 0 | 0 |
| *Klebsiella pneumoniae* (VA13414) | 257 | 234 | 1 | 0 |
| *Klebsiella pneumoniae* (BK12739) | 244 | 243 | 0 | 0 |
| *Klebsiella pneumoniae* (VA18342) | 247 | 230 | 0 | 0 |
| *Klebsiella pneumoniae* (VA23130) | 111 | 99 | 3 | 3 |
| *Klebsiella pneumoniae* (BK13728) | 53 | 55 | 0 | 1 |
| *Klebsiella pneumoniae* (UR14350) | 14 | 5 | 0 | 0 |
| *Klebsiella pneumoniae* (D411554) | 11 | 12 | 3 | 3 |
| *Klebsiella pneumoniae* (VA13324) | 13 | 9 | 0 | 1 |
| *Klebsiella pneumoniae* (TP3419) | 14 | 5 | 12 | 17 |
| *Klebsiella aerogenes* (BC05_run91) | 22 | 15 | 1 | 1 |
| *Shewanella algae* (BC11_run90) | 40 | 34 | 0 | 0 |
| *Acinetobacter soli* (BC11_run91) | 288 | 239 | 0 | 0 |
| *Acinetobacter junii* (BC12_run91) | 42 | 44 | 1 | 2 |
| *Enterobacter hormaechei* (BC21_run89) | 345 | 333 | 79 | 69 |
| *Citrobacter freundii* (BC22_run90) | 22 | 29 | 57 | 64 |

**Supplementary Table 3:** Evaluation of errors in reads for different species, prepared with predecessor sequencing kit SQK-LSK109. Only the ambiguous positions on the chromosomes are considered in this table. The evaluation was done using MPOA workflow.

| Species | N(ATCG) | W(AT) | S(CG) | M(AC) | K(TG) | R(AG) | Y(TC) |
|---|---|---|---|---|---|---|---|
| *Achromobacter bronchisepticus* | 408 | 0 | 0 | 0 | 0 | 4 | 4 |
| *Acinetobacter baumannii* | 92 | 1 | 2 | 1 | 1 | 63 | 50 |
| *Acinetobacter baumannii* | 601 | 1 | 0 | 0 | 0 | 42 | 53 |
| *Acinetobacter baumannii* | 23 | 3 | 0 | 0 | 1 | 36 | 27 |
| *Acinetobacter baumannii* | 10381 | 0 | 0 | 0 | 1 | 3 | 4 |
| *Acinetobacter baumannii* | 1615 | 0 | 0 | 0 | 0 | 39 | 36 |
| *Acinetobacter baumannii* | 1936 | 0 | 0 | 0 | 0 | 1 | 1 |
| *Acinetobacter baumannii* | 132 | 1 | 0 | 0 | 0 | 3 | 2 |
| *Acinetobacter baumannii* | 3569 | 0 | 0 | 0 | 0 | 0 | 2 |
| *Chryseobacterium bernardetii_A* | 18413 | 19 | 6 | 8 | 13 | 27 | 26 |
| *Chryseobacterium indologenes* | 14 | 2 | 0 | 0 | 1 | 20 | 16 |
| *Elizabethkingia anophelis* | 29 | 2 | 0 | 3 | 2 | 7 | 1 |
| *Elizabethkingia anophelis* | 136 | 5 | 0 | 0 | 0 | 4 | 5 |
| *Enterobacter asburiae* | 10 | 0 | 0 | 0 | 0 | 4 | 1 |
| *Enterobacter cloacae* | 5 | 0 | 0 | 0 | 1 | 83 | 55 |
| *Enterobacter ludwigii* | 2 | 0 | 0 | 0 | 0 | 0 | 3 |
| *Escherichia coli* | 45 | 0 | 0 | 0 | 0 | 2 | 3 |
| *Escherichia coli* | 67 | 15 | 33 | 17 | 15 | 50 | 47 |
| *Escherichia coli* | 19 | 0 | 0 | 0 | 0 | 27 | 34 |
| *Escherichia coli* | 24736 | 0 | 0 | 0 | 0 | 0 | 1 |
| *Escherichia flexneri* | 10 | 1 | 0 | 0 | 0 | 20 | 32 |
| *Klebsiella pneumoniae* | 0 | 0 | 0 | 0 | 0 | 2 | 1 |
| *Klebsiella pneumoniae* | 13 | 1 | 0 | 0 | 0 | 27 | 20 |
| *Klebsiella pneumoniae* | 4 | 1 | 0 | 0 | 0 | 7 | 4 |
| *Klebsiella pneumoniae* | 6974 | 0 | 0 | 0 | 1 | 7 | 5 |
| *Klebsiella quasipneumoniae* | 1210 | 0 | 0 | 0 | 2 | 1 | 3 |
| *Pseudomonas aeruginosa* | 18871 | 0 | 0 | 0 | 0 | 28 | 18 |
| *Pseudomonas aeruginosa* | 958 | 1 | 51 | 2 | 11 | 35 | 30 |
| *Pseudomonas aeruginosa* | 41 | 0 | 10 | 0 | 5 | 12 | 12 |
| *Pseudomonas aeruginosa* | 21 | 1 | 8 | 4 | 0 | 34 | 38 |

| Species | N(ATCG) | W(AT) | S(CG) | M(AC) | K(TG) | R(AG) | Y(TC) |
|---|---|---|---|---|---|---|---|
| *Pseudomonas aeruginosa* | 16022 | 0 | 1 | 0 | 0 | 19 | 34 |
| *Pseudomonas aeruginosa* | 15763 | 0 | 0 | 0 | 2 | 14 | 5 |
| *Pseudomonas aeruginosa* | 28 | 0 | 0 | 0 | 0 | 2 | 2 |
| *Pseudomonas aeruginosa* | 7612 | 0 | 0 | 1 | 6 | 12 | 13 |
| *Pseudomonas aeruginosa* | 10 | 0 | 0 | 1 | 0 | 11 | 13 |
| *Pseudomonas aeruginosa* | 129 | 0 | 0 | 1 | 0 | 10 | 6 |
| *Salmonella enterica* | 312 | 0 | 0 | 0 | 0 | 2 | 1 |
| *Serratia marcescens* | 31 | 0 | 0 | 0 | 0 | 15 | 17 |
| *Serratia marcescens* | 596 | 0 | 0 | 0 | 0 | 5 | 4 |
| *Serratia marcescens* | 43 | 1 | 0 | 0 | 1 | 7 | 8 |
| *Serratia marcescens* | 3 | 0 | 0 | 0 | 0 | 1 | 9 |

**Supplementary Table 4:** Comparison of errors in reads for 11 different species when basecalled with different models and basecaller. Only the most problematic observed ambiguous positions, R and Y, within the chromosome, are included in this table. The evaluation was done using MPOA workflow.

| | Guppy (*dna_r10.4.1_e8.2_260bps_sup.cfg*) | | Dorado (*res_dna_r10.4.1_e8.2_400bps_sup.cfg*) | |
|---|---|---|---|---|
| Species | R(AG) | Y(TC) | R(AG) | Y(TC) |
| *Pseudomonas aeruginosa* | 49 | 54 | 15 | 14 |
| *Pseudomonas aeruginosa* | 1091 | 1189 | 896 | 791 |
| *Pseudomonas aeruginosa* | 1 | 1 | 1 | 0 |
| *Pseudomonas aeruginosa* | 138 | 100 | 37 | 38 |
| *Pseudomonas aeruginosa* | 0 | 1 | 0 | 0 |
| *Pseudomonas aeruginosa* | 138 | 136 | 61 | 51 |
| *Pseudomonas aeruginosa* | 0 | 0 | 0 | 0 |
| *Pseudomonas aeruginosa* | 33 | 40 | 9 | 14 |
| *Pseudomonas aeruginosa* | 0 | 0 | 0 | 0 |
| *Pseudomonas aeruginosa* | 111 | 99 | 52 | 48 |
| *Pseudomonas aeruginosa* | 33 | 40 | 9 | 14 |
| *Klebsiella pneumoniae* | 47 | 42 | 17 | 22 |

| | | | | |
|---|---|---|---|---|
| *Klebsiella pneumoniae* | 54 | 40 | 22 | 12 |
| *Acinetobacter baumannii* | 1 | 0 | 0 | 0 |
| *Acinetobacter baumannii* | 10 | 9 | 2 | 1 |
| *Achromobacter xylosoxidans* | 6 | 4 | 1 | 0 |
| *Citrobacter freundii* | 48 | 38 | 5 | 10 |
| *Enterobacter hormaechei* | 4 | 6 | 3 | 1 |
| *Escherichia flexneri* | 85 | 86 | 32 | 29 |
| *Klebsiella_A oxytoca* | 5 | 9 | 1 | 1 |
| *Klebsiella_A michiganensis* | 22 | 29 | 18 | 19 |
| *Salmonella enterica* | 82 | 85 | 23 | 19 |
| *Stenotrophomonas maltophilia* | 228 | 167 | 198 | 170 |

### Supplementary Figures

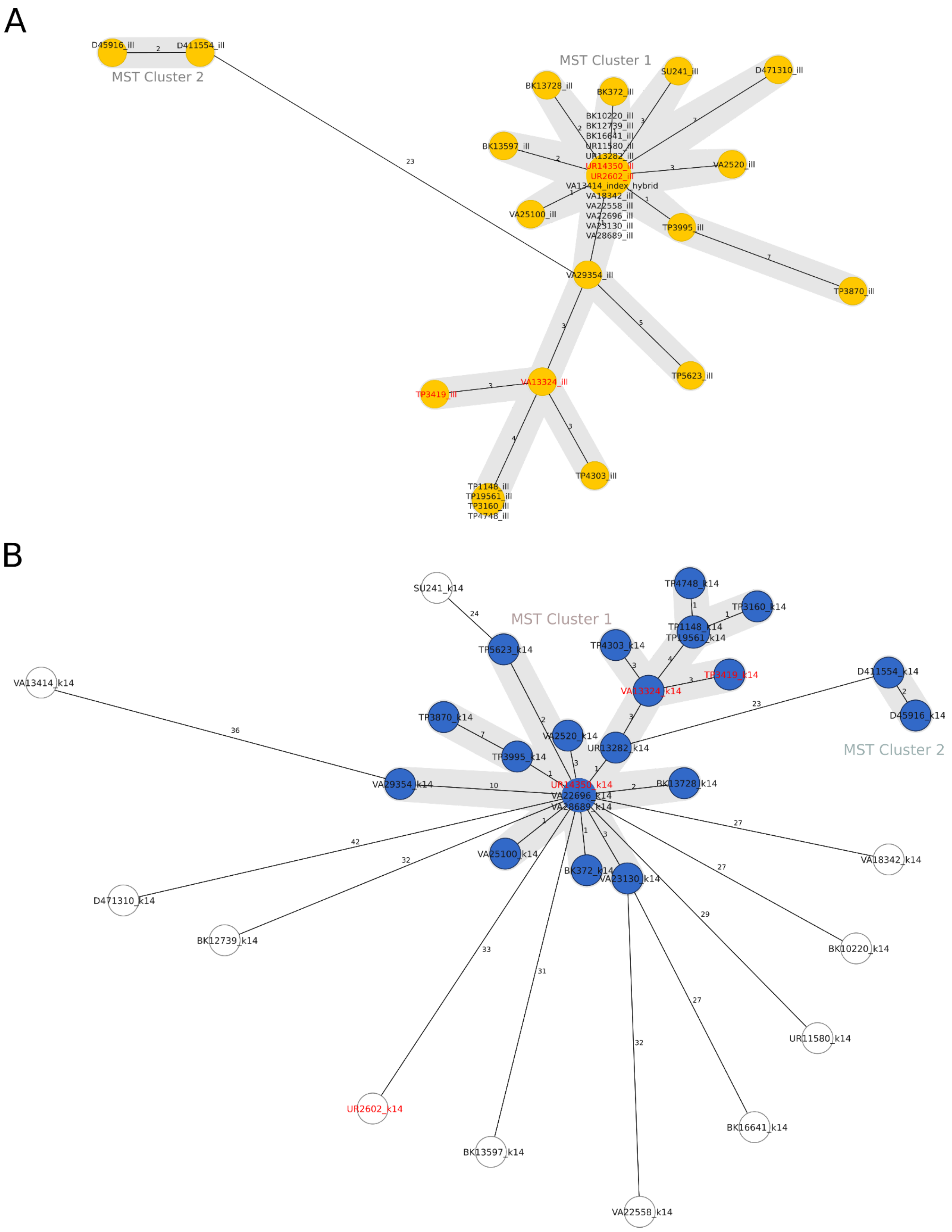


**Supplementary Figure 1:** Both Minimal spanning trees (pairwise ignore missing values showing) based on 2358 genes of each 33 *K. pneumoniae* outbreak samples. Samples used in the manuscript for Figure 1 are highlighted in red. Samples with allelic differences ≤15 are considered as part of the cluster. **A:** Assemblies built from Illumina sequencing data (yellow). **B:** Assemblies built from Nanopore Kit14 sequencing data (blue). Outlier samples not in the outbreak cluster and showing significant differences from the Illumina data are shown in white. Nodes (samples) are connected by lines depicting the distance by numbers of allelic differences. Loci are considered different if one or more bases change between the samples. Loci without allelic differences are described as being the same.

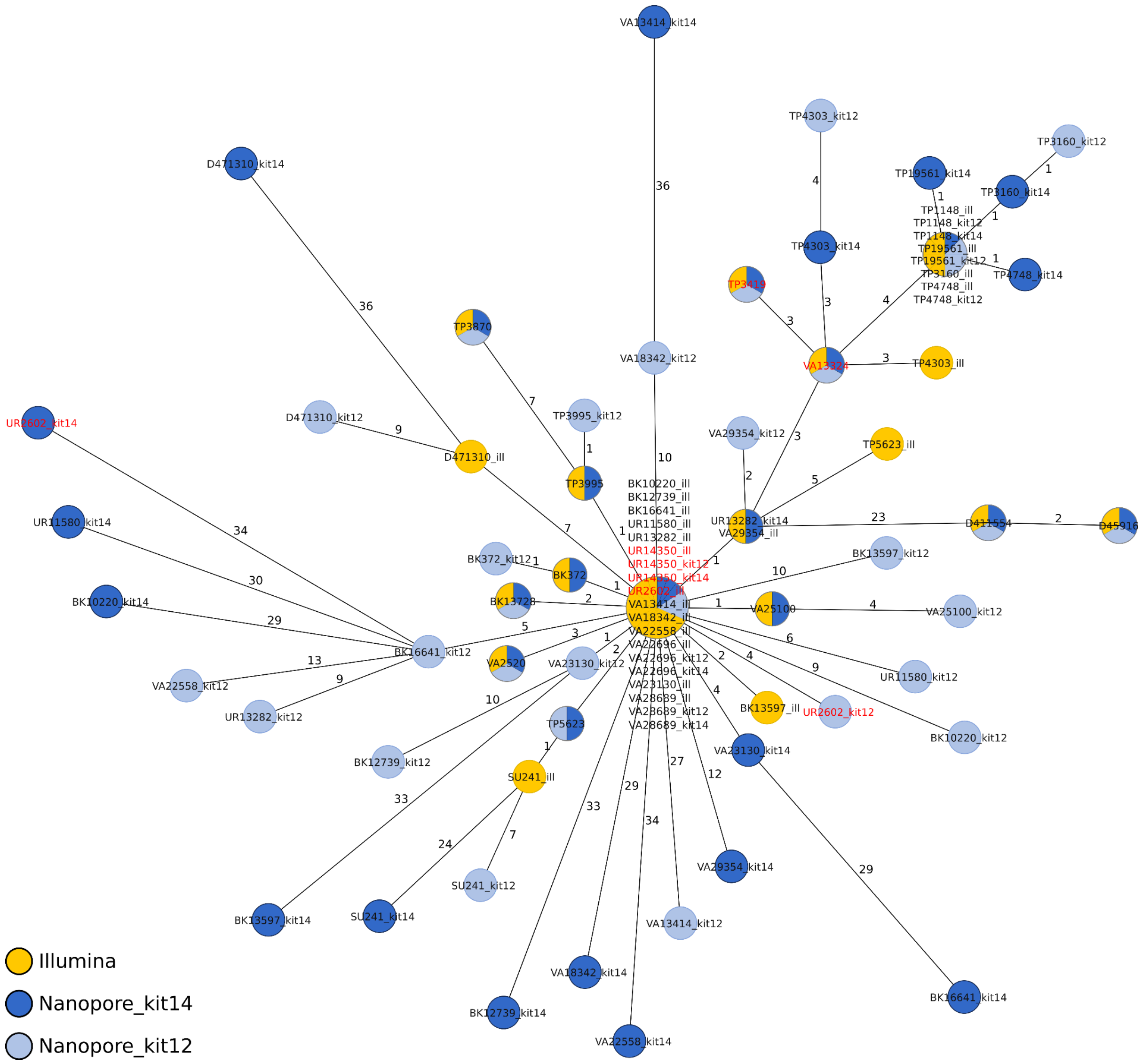


**Supplementary Figure 2:** Minimum spanning trees (pairwise ignore missing values showing) of each 33 *K. pneumoniae* outbreak samples based on 2358 genes to compare the allelic variations between Illumina genomes, Nanopore SQK-NBD114.24 (kit14) and SQK-NBD114.24 (kit12). Samples used in the manuscript for Figure 1 are highlighted in red. in Nodes (samples) are connected by lines depicting the distance by numbers of allelic differences. Loci are considered different if one or more bases change between the samples. Loci without allelic differences are described as being the same. Nodes are colored according to the sequencing technology used. Samples with allelic differences ≤15 are considered as part of the cluster.

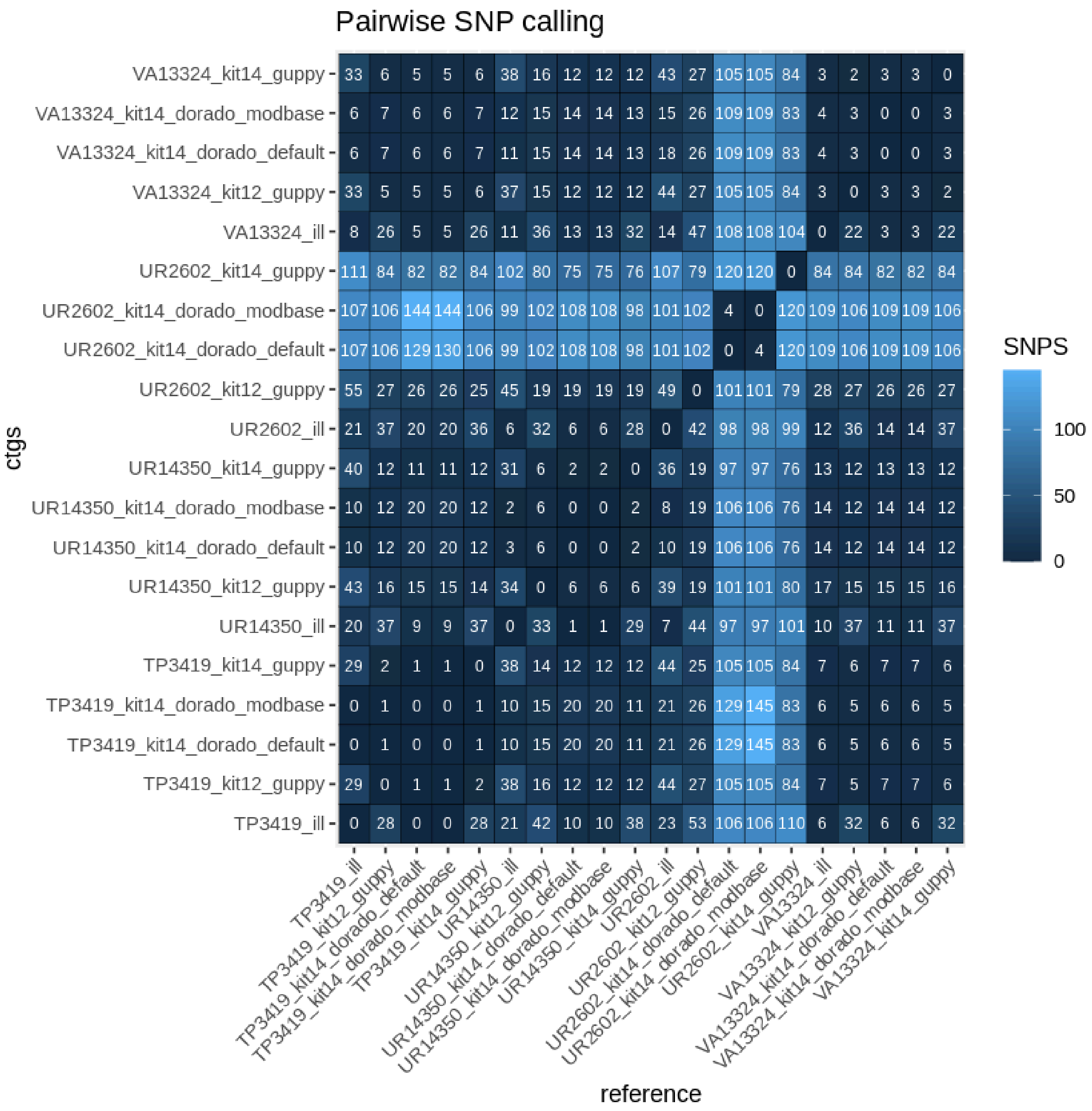


**Supplementary Figure 3:** Heatmap of pairwise SNP comparison of four genomes based on Snippy analysis. The four isolates used were each prepared with Illumina (gold standard and reference), Kit 14, and Kit 12 and basecalled with each respective Guppy “super accurate” basecalling model (see methods “Basecalling and Assembly”). All Kit 14-prepared isolates were additionally basecalled with Dorado using the default and a modification-aware model.

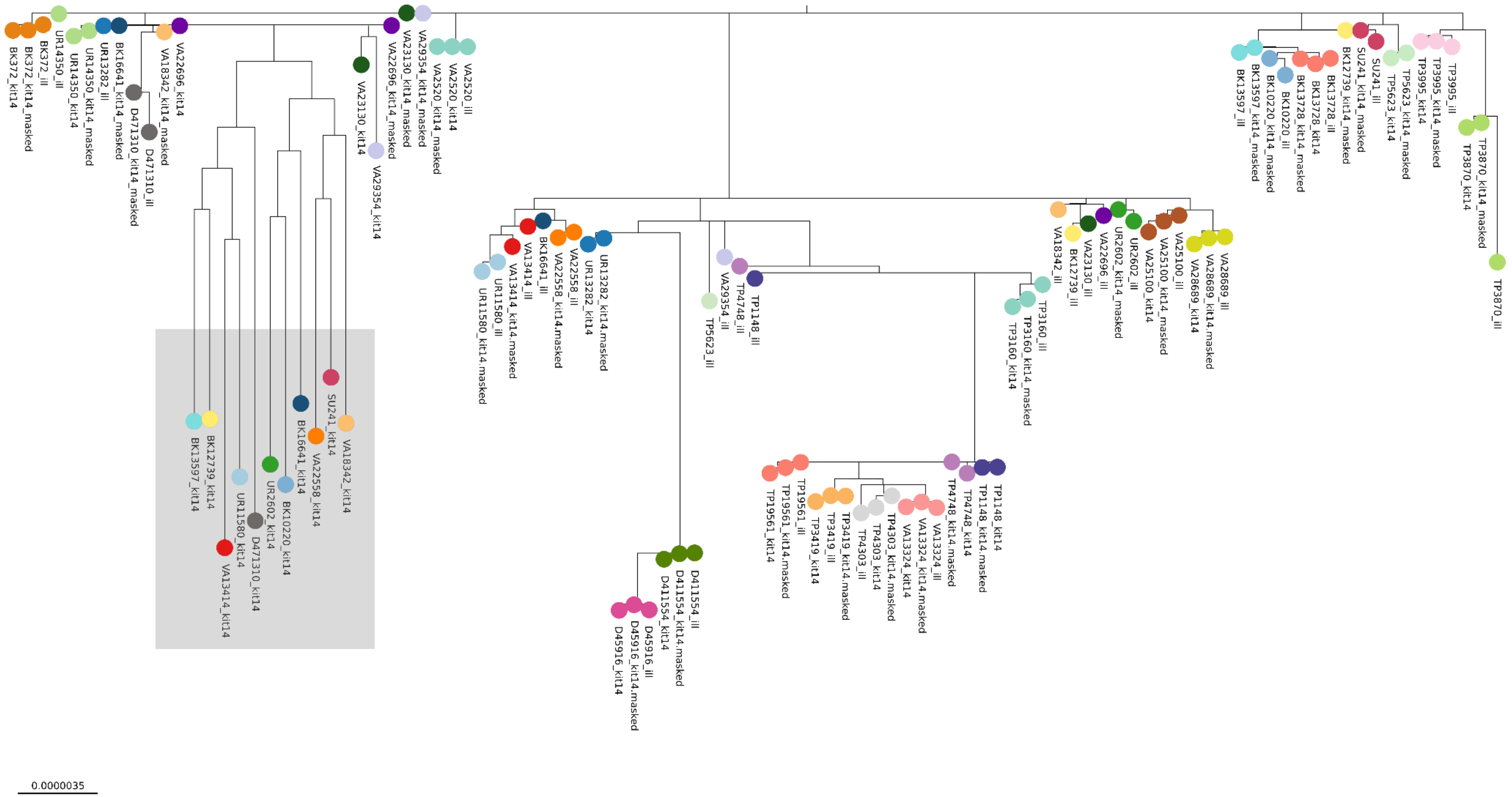


**Supplementary Figure 4:** Phylogenetic tree based on core genome SNP alignment to figure the genetic distances between 33 *K. pneumoniae* outbreak samples (colored nodes), sequenced using Illumina (ill) and Nanopore SQK-NBD114.24 (kit14) compared to the masked Kit 14 assemblies (masked). Detected outliers are highlighted in grey.

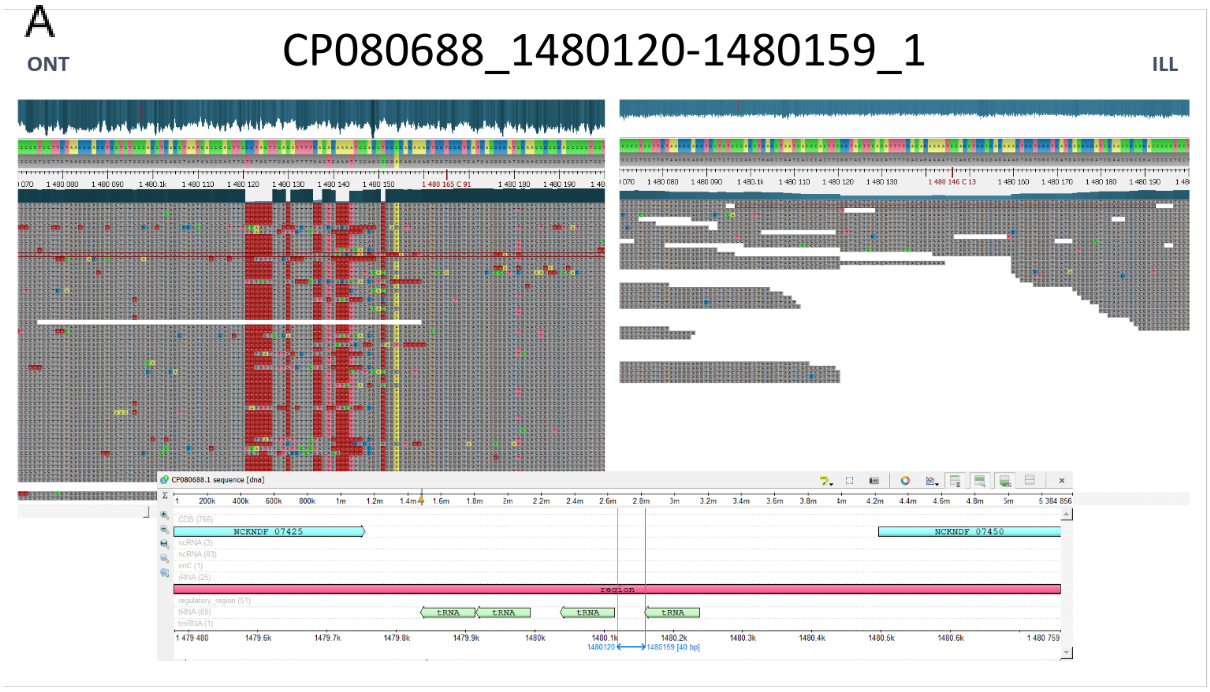


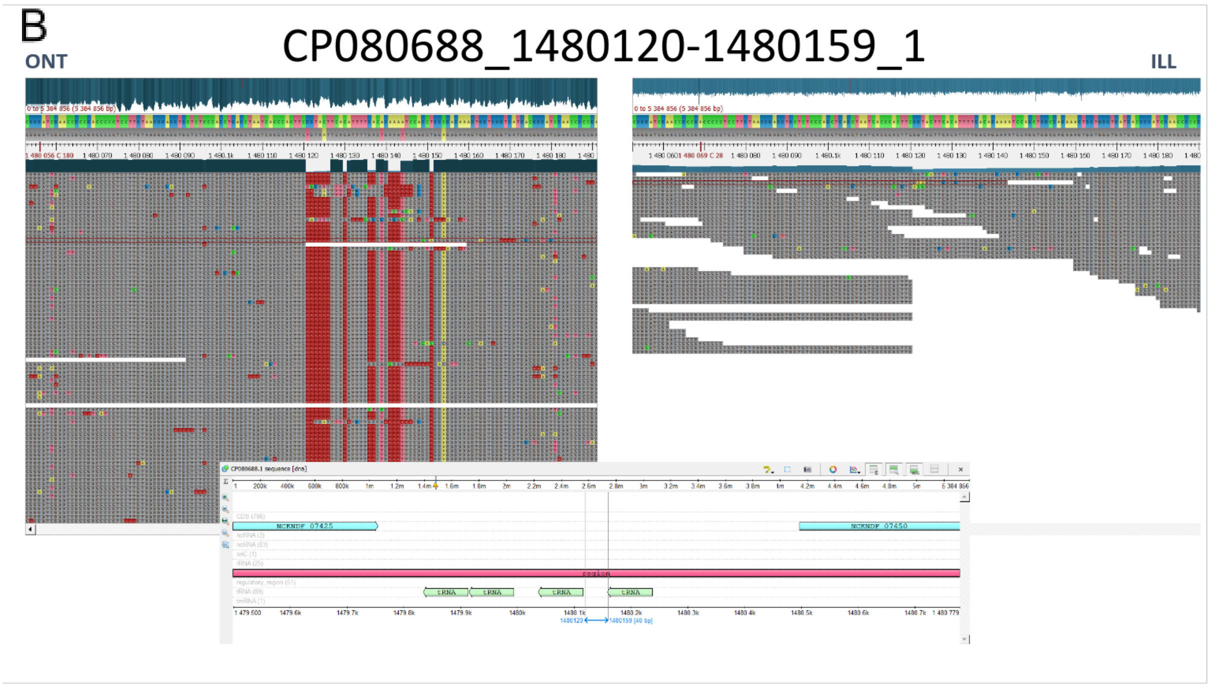


**Supplementary Figure 5:** Identified errors in Illumina Assemblies by mapped reads of the same sample and sequence region Nanopore (left) and Illumina (right) against Illumina Hybrid Assembly of Index patients in the position 1480120-1480159 on the chromosome. **A:** VA28689 **B**: BK16641.

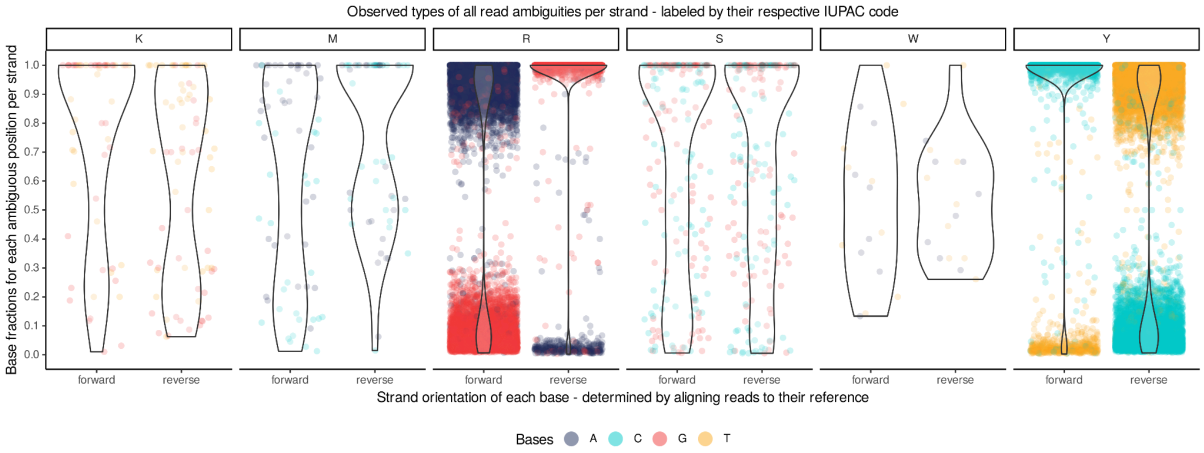


**Supplementary Figure 6:** Violin chart showing the ratio between two bases within the mapped read data distinguished by strand orientation for 19 *Pseudomonas aeruginosa* samples.

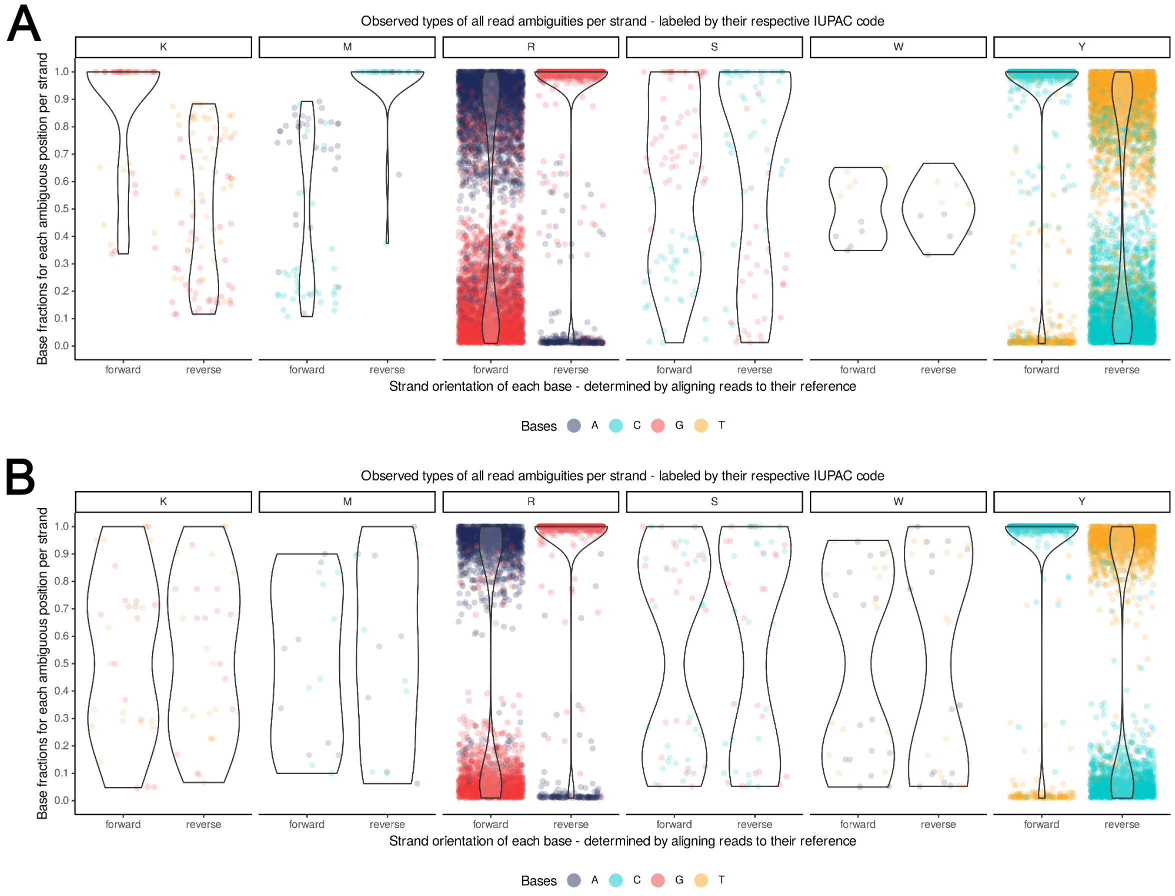


**Supplementary Figure 7: Mapping strategies - minimap2 vs bwa** Frequency plot over minimap2 (A) and bwa (B) mapped reads on polished assembly.

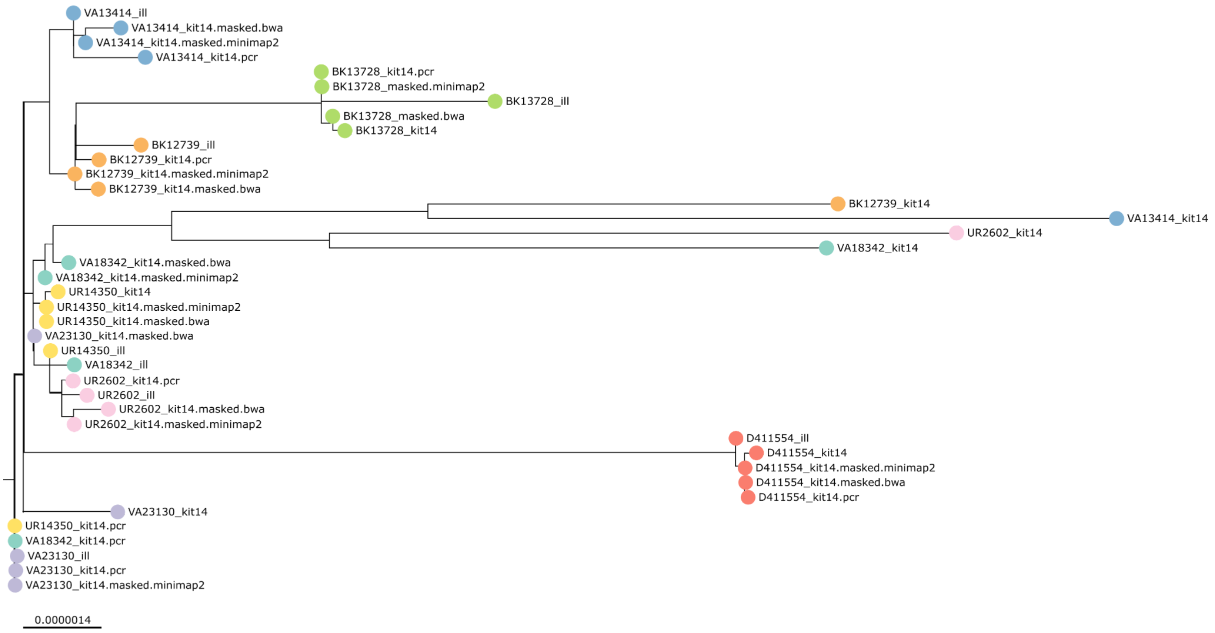


**Supplementary Figure 8: Mapping strategies - minimap2 vs BWA** Phylogenetic tree based on core genome SNP alignment to figure the genetic distances between eight *K. pneumoniae* outbreak samples (colored nodes), prepared with Illumina (ill), Nanopore SQK-NBD114.24 (kit14) and SQK-RPB114.24 (pcr) compared to masked Kit 14 assemblies based on minimap2 mapping (masked.minimap2) and masked Kit 14 assemblies based on BWA mapping (masked.bwa).

### Supplementary Code

**Supplementary Code 1:** Used Code for Assembly, Polishing and phylogenetic Analysis

```
flye --plasmids --meta -t ${task.cpus} --nano-hq ${read} -o
assembly
minimap2 -x map-ont -t ${task.cpus} ${assembly} ${read} >
${name}.paf
minimap2 -ax map-ont ${assembly} ${read} | samtools view -bS - |
samtools sort -@ ${task.cpus} - > ${name}_${technology}_sorted.bam
racon -t ${task.cpus} ${read} ${mapping} ${assembly} >
${name}_consensus.fasta
medaka_consensus -i ${read} -d ${consensus} -o polished -t
${task.cpus} -m \${POLISHMODEL}
FastTree -gtr -nt ${clean_core_alignment} > clean.core.tree.nwk
for fasta in genomes/*
      do
                FILENAME=\$(basename \${fasta%.*})
                snippy --cpus ${task.cpus} --ram ${maxmemory}
--outdir ${cluster_ID}/\${FILENAME}/ --ref reference/* --ctgs
\${fasta}
           done
snippy-core --ref reference/* --prefix results ${cluster_ID}/*/
```
