## Supplementary Code 2 for "Accurate bacterial outbreak tracing with Oxford Nanopore sequencing and reduction of methylation-induced errors": create_logo.R

```
#!/usr/bin/env Rscript

# Load the required packages
require(ggplot2)
require(ggseqlogo)
require(readr)

# Inputs
INPUTS = commandArgs(trailingOnly=TRUE)

samplename=INPUTS[1]

if (!file.exists("W.regions.fasta")) { Wpos <- list() } else { Wpos <- read_lines("W.regions.fasta", skip_empty_rows = TRUE) }
if (!file.exists("S.regions.fasta")) { Spos <- list() } else { Spos <- read_lines("S.regions.fasta", skip_empty_rows = TRUE) }
if (!file.exists("M.regions.fasta")) { Mpos <- list() } else { Mpos <- read_lines("M.regions.fasta", skip_empty_rows = TRUE) }
if (!file.exists("K.regions.fasta")) { Kpos <- list() } else { Kpos <- read_lines("K.regions.fasta", skip_empty_rows = TRUE) }
if (!file.exists("R.regions.fasta")) { Rpos <- list() } else { Rpos <- read_lines("R.regions.fasta", skip_empty_rows = TRUE) }
if (!file.exists("Y.regions.fasta")) { Ypos <- list() } else { Ypos <- read_lines("Y.regions.fasta", skip_empty_rows = TRUE) }
if (!file.exists("B.regions.fasta")) { Bpos <- list() } else { Bpos <- read_lines("B.regions.fasta", skip_empty_rows = TRUE) }
if (!file.exists("D.regions.fasta")) { Dpos <- list() } else { Dpos <- read_lines("D.regions.fasta", skip_empty_rows = TRUE) }
if (!file.exists("H.regions.fasta")) { Hpos <- list() } else { Hpos <- read_lines("H.regions.fasta", skip_empty_rows = TRUE) }
if (!file.exists("V.regions.fasta")) { Vpos <- list() } else { Vpos <- read_lines("V.regions.fasta", skip_empty_rows = TRUE) }

# combining and dropping fasta headers
combined = list(Wpos, Spos, Mpos, Kpos, Rpos, Ypos, Bpos, Dpos, Hpos, Vpos)
combined <- lapply(combined, function(x){x[!grepl(">",x)]})
combined <- lapply(combined, function(x) c(paste0(length(x), ' masked regions for '), x))
#renaming
name_list <- sapply(combined, "[[", 1)
name_list2 <- c("W(AT)","S(CG)","M(AC)","K(TG)","R(AG)","Y(TC)","B(TCG)","D(ATG)","H(ATC)","V(ACG)")
								
name_list_comb <-paste0(name_list,name_list2)
names(combined) <- name_list_comb
combined <- lapply(combined, function(x) x[-1])

# removes empty lines
combined <-  combined[sapply(combined, length) > 0]

# WSMKRYBDHV
cs_gaps <-
  make_col_scheme(
    chars = c("A", "C", "T", "G", "W", "S", "M", "K", "R", "Y", "B", "D", "H", "V", "N"),
    cols = c("#588157", "#B3001B", "#2E4A76", "#E08E45", "#433347", "#433347", "#433347", 
             "#433347", "#433347", "#433347", "#433347", "#433347", "#433347", "#433347", "#433347")
  )

plot <- ggseqlogo(combined, method = 'bits', namespace = cs_gaps$letter, col_scheme = cs_gaps, ncol = 1) + 
        labs(title = samplename) 


svg("logo.svg", height=as.numeric(length(combined))*2)
plot
dev.off()
```
