## Supplementary Code 2 for "Accurate bacterial outbreak tracing with Oxford Nanopore sequencing and reduction of methylation-induced errors": violin_chart.R

```
#!/usr/bin/env Rscript

library(ggplot2)
require(readr)

names <- c('orientation', 'deg', 'Bases', 'ratio')
df1 <- read_tsv("input.tsv", col_names=names)

plot <- ggplot(df1, aes(x=orientation, y=ratio)) +
  scale_fill_brewer(palette="Accent") +
  geom_point( aes(x=orientation, y=ratio, colour=Bases), shape = 16,size=2, position = position_jitterdodge(jitter.width = 0.8, jitter.height = 0, dodge.width = 0), alpha=0.2, show.legend = TRUE) +
  scale_color_manual(values = c("#24305E", "#00CCCC", "#f43c3c", "#FBAC23")) +   # alternative #5C5C5C
  coord_cartesian(ylim=c(0,1)) +
  scale_y_continuous(breaks = seq(0,1, by=0.1)) +
  geom_violin(alpha=0.2, position = position_dodge(width = .75)) +
  facet_grid(cols = vars(deg), scales = "free", space = "free") +
  ylab(  c("Base fractions for each ambiguous position per strand")  ) +
  xlab(  c("Strand orientation of each base - determined by aligning reads to their reference")  ) +
  theme_classic() +
  theme(legend.position="bottom", plot.subtitle = element_text(hjust = 0.5))  +
  labs(subtitle = "Observed types of all read ambiguities per strand - labeled by their respective IUPAC code") +
  guides(colour = guide_legend(override.aes = list(size=5, alpha=0.5)))


svg("frequency.svg", height=5, width= (as.numeric(length(unique(df1$deg))) * 2) + 1)  # , *2   length(unique(df$col))
plot
dev.off()
```
