## Supplementary Code 2 for "Accurate bacterial outbreak tracing with Oxford Nanopore sequencing and reduction of methylation-induced errors": main.nf

```
#!/usr/bin/env nextflow
nextflow.enable.dsl=2


/*
* Nextflow -- mask regions for outbreak analysis
* Author:
*/


/************************** 
* Help messages & user inputs & checks
**************************/
// help message
    if (params.help) { exit 0, helpMSG() }


// error codes
    if (params.profile) {
        exit 1, "--profile is WRONG use -profile" }

    if ( workflow.profile == 'standard' ) { 
        "NO EXECUTION PROFILE SELECTED, using [-profile local,docker]" }


    if ( !params.fastq &&  !params.fasta ) {
        exit 1, "input missing, use [--fasta] and [--fastq]"}

 
// nanopore reads
    if (params.fastq) { nano_input_ch = Channel
            .fromPath( params.fastq, checkIfExists: true)
            .map { file -> tuple(file.simpleName, file) }
            }

    if (params.fasta) { fasta_input_ch = Channel
            .fromPath( params.fasta, checkIfExists: true)
            .map { file -> tuple(file.simpleName, file) }
            }


/************************** 
* WORKFLOWS
**************************/

include { mask_regions_wf } from './workflows/mask_regions.nf' 
include { identify_logo_wf } from './workflows/identify_logo.nf'

/************************** 
* MAIN WORKFLOW 
**************************/

workflow {
    defaultMSG()
    if (params.keep_coordinates) {keepMSG()}
   
        identify_logo_wf(mask_regions_wf(nano_input_ch, fasta_input_ch)) 
             
}


/*************  
* LOG INFO DEFINITIONS
*************/

def helpMSG() {
    c_green = "\033[0;32m";
    c_reset = "\033[0m";
    c_yellow = "\033[0;33m";
    c_blue = "\033[0;34m";
    c_dim = "\033[2m";
    log.info """
    ____________________________________________________________________________________________
    
    Workflow: MPOA - Mask bases in Pathogens for Outbreak Analysis

    ${c_yellow}Usage example:${c_reset}
    nextflow run replikation/MPOA --fastq '*.fastq' --fasta '*.fasta' -profile local,docker

    ${c_yellow}Inputs (Mandatory):
     ${c_green}--fastq ${c_reset}        e.g.: 'sample1.fastq' or '*.fastq' or '*/*.fastq'
     ${c_green}--fasta ${c_reset}        e.g.: 'sample1.fasta' or '*.fasta' 
    Reads and Genome files are matched based on the first word before the first dot in their filename.
      Matches: ${c_green}Sample1${c_reset}.clean.fasta ${c_green}Sample1${c_reset}.fastq.gz
      This NOT: ${c_yellow}clean${c_reset}.Sample1.clean.fasta ${c_yellow}Sample1${c_reset}.fastq.gz

    ${c_yellow}Workflow settings${c_reset}  
     ${c_blue}--frequency ${c_reset}    Turns on frequency calculation of bases [default: $params.frequency]
     ${c_blue}--depth X ${c_reset}      Masks regions with a sequencing depth below X with N's [default: $params.depth]
     ${c_blue}--motif X ${c_reset}      Upstream and Downstream length of sequence Motif [default: $params.motif]
     ${c_blue}--mapper  ${c_reset}      Read mapper to use: minimap2, bwa [default: $params.mapper]

    ${c_yellow}Options (optional)${c_reset}
     --keep_coordinates     Output all reference positions, including low coverage regions.
                            Ignore insertions in the consensus and mask deletions with "*".
                            This will add '-aa --show-ins no --show-del no' to samtools consensus [default: $params.keep_coordinates]
     --cores                Amount of cores for a process (local use) [default: $params.cores]
     --max_cores            Max amount of cores for poreCov to use (local use) [default: $params.max_cores]
     --memory               Available memory [default: $params.memory]
     --output               Name of the result folder [default: $params.output]
     --workdir              Defines the path to the temporary files [default: $params.workdir]
     --cachedir             Defines the path where singularity images are cached [default: $params.cachedir]

    ${c_yellow}Execution/Engine profiles (choose executer and engine)${c_reset}
    MPOA supports profiles to run via different ${c_green}Executers${c_reset} and ${c_blue}Engines${c_reset} 
    Examples:
     -profile ${c_green}local${c_reset},${c_blue}docker${c_reset}
     -profile ${c_green}slurm${c_reset},${c_blue}singularity${c_reset}

      ${c_green}Executer${c_reset} (choose one):
       local
       slurm
      ${c_blue}Engines${c_reset} (choose one):
       docker
       singularity

    Note: The singularity profile automatically passes the following environment variables to the container. 
    to ensure execution on HPCs: HTTPS_PROXY, HTTP_PROXY, http_proxy, https_proxy, FTP_PROXY, ftp_proxy
    """.stripIndent()
}

def defaultMSG() {
    c_green = "\033[0;32m";
    c_reset = "\033[0m";
    c_yellow = "\033[0;33m";
    c_blue = "\033[0;34m";
    c_dim = "\033[2m";
    log.info """
    MPOA
    \u001B[1;30m______________________________________\033[0m

    \u001B[36mMask Pathogens for Outbreak Analysis\033[0m
    \u001B[1;30m______________________________________\033[0m
    Profile:                $workflow.profile
    Current User:           $workflow.userName
    Nextflow-version:       $nextflow.version
    Starting time:          $nextflow.timestamp

    Frequency calculation:
        --keep_coordinates  $params.keep_coordinates
        --frequency         $params.frequency
        --mapper            $params.mapper


        --workdir           $params.workdir
        --output            $params.output
        --max_cores         $params.max_cores        
        --cores             $params.cores
        --mem               $params.memory
    \u001B[1;30m______________________________________\033[0m
    """.stripIndent()
}

def keepMSG() {
    c_green = "\033[0;32m";
    c_reset = "\033[0m";
    c_yellow = "\033[0;33m";
    c_blue = "\033[0;34m";
    c_dim = "\033[2m";
    log.info """
    \u001B[36mKeep coordinates option is activated!\033[0m
    \u001B[1;30m______________________________________\033[0m
    ATTENTION: The --keep_coordinates parameter will activate '-aa --show-ins no --show-del no' in the samtools consensus command. 

    ${c_yellow}This will cause all reference positions to appear in the consensus regardless of their sequence coverage. 

    Insertions will not be reported in the consensus to keep the original coordinates.

    Deletions will not be deleted in the consensus but masked by "*". 

    When using this parameter, you can keep the coordinates of your input reference FASTA intact.
    This is helpful if you are searching problematic sites with respect to your input unmasked reference sequence. 

    But be aware that you might not directly use the output masked FASTA in downstream applications. 
    \u001B[1;30m______________________________________\033[0m
    """.stripIndent()
}
```
