## Supplementary Code 2 for "Accurate bacterial outbreak tracing with Oxford Nanopore sequencing and reduction of methylation-induced errors": README.md

<p align="center">
  <img src="data/logo/mobile_logo.png" width="800" title="Workflow">
</p>

**MPOA | Workflow to mask pathogens for outbreak analysis**   
===
![](https://img.shields.io/github/v/release/replikation/MPOA)
![](https://img.shields.io/badge/uses-Docker-blue.svg)
![](https://img.shields.io/badge/licence-GPL--3.0-lightgrey.svg)


[![Twitter Follow](https://img.shields.io/twitter/follow/maralohde.svg?style=social)](https://twitter.com/maralohde) 
[![Twitter Follow](https://img.shields.io/twitter/follow/gcloudChris.svg?style=social)](https://twitter.com/gcloudChris) 

**Citation:**
>Nanopore sequencing for accurate bacterial outbreak tracing <br>
>Mara Lohde, Gabriel E. Wagner, Johanna Dabernig-Heinz, Adrian Viehweger, Sascha D. Braun, Stefan Monecke,Celia Diezel, Claudia Stein, Mike Marquet, Ralf Ehricht, Mathias W. Pletz, Christian Brandt <br>
>https://www.biorxiv.org/content/10.1101/2023.09.15.556300v2

## What is this Repo?
* A quick workflow to mask fasta files for outbreak analysis
* It mitigates false base calls from e.g., modified bases by masking all such positions that are "uncertain"
* Figures and overviews are provided to assess weather your samples might be affected by such an issue

<p align="center">
  <img src="data/figures/MPOA_flowchart.png" width="700" title="Workflow">
</p>

## Documentation 
https://maralohde.github.io/MPOA-documentation/

# Quick installation
## 1.1 Nextflow (the workflow manager)
* MPOA needs [Nextflow](https://www.nextflow.io/index.html) and java run time (default-jre)
    * install java run time via:  `sudo apt install -y default-jre`
    * install Nextflow e.g.:  `curl -s https://get.nextflow.io | bash && sudo mv nextflow /bin && sudo chmod 770 /bin/nextflow`
## 1.2 Docker
* Installation [here](https://docs.docker.com/v17.09/engine/installation/linux/docker-ce/ubuntu/#install-docker-ce) (recommended), alternatively via: `sudo apt install -y docker`
* Add Docker to the user: `sudo usermod -a -G docker $USER`
## 1.3 Singularity
* As an alternative to Docker you can also install and use Singularity, e.g. on a HPC
* Installation [here](https://apptainer.org/docs/)
* Note, that with Singularity the following environment variables are automatically passed to the container to ensure execution on HPCs: `HTTPS_PROXY`, `HTTP_PROXY`, `http_proxy`, `https_proxy`, `FTP_PROXY` and `ftp_proxy`.

# Quick start
* Reads and genomes are matched by their name before the first "."
* E.g. genome file **sample1**.test.fasta matches with read file **sample1**.fastq.gz

```bash
#install
nextflow pull replikation/MPOA

#help
nextflow run replikation/MPOA -profile local,docker --help

#example run
nextflow run replikation/MPOA --fastq '*.fastq' --fasta '*.fasta' -profile local,docker

#example run on a HPC
nextflow run replikation/MPOA --fastq '*.fastq' --fasta '*.fasta' -profile slurm,singularity
```
