## Supplementary Code 2 for "Accurate bacterial outbreak tracing with Oxford Nanopore sequencing and reduction of methylation-induced errors": identify_logo.nf

```
include { count_positions } from './process/count_positions.nf'
include { extract_sequence } from './process/extract_sequence.nf'
include { plot_motifs } from './process/plot_motifs.nf'
include { get_frequency } from './process/get_frequency.nf'
include { plot_frequency} from './process/plot_frequency.nf'

workflow identify_logo_wf {
    take:   
        fasta // masked and depth masked
        bam // masked bam (not dm masked)
    main: 

        // get positions
        extract_sequence(count_positions(fasta))

        // visualize motifs
        plot_motifs(extract_sequence.out)

        // plot frequency per deg. base
        if (params.frequency) {
            get_frequency(
                count_positions.out // val path( fastadm and masked), path (positions deg bases)
                .join(bam)
            )

            // collect all frequency results
            plot_frequency(get_frequency.out.map{it -> it[1]}.collect())
        }      

    emit:  extract_sequence.out
}
```
