## Supplementary Code 2 for "Accurate bacterial outbreak tracing with Oxford Nanopore sequencing and reduction of methylation-induced errors": mask_regions.nf

```
include { minimap2 } from './process/minimap2.nf'
include { plasflow; plasflow_degen } from './process/plasflow.nf'
include { bedtools } from './process/bedtools.nf'
include { bwa } from './process/bwa.nf'


workflow mask_regions_wf {
    take:   
        fastq
        fasta
    main: 

    // join channels
    combined_ch = fasta.join(fastq, by:0)

    // check if everything is alright
    no_match = combined_ch.ifEmpty{ log.info "\033[0;33mCould not match any reads to genomes, please read the help via --help\033[0m" }

    if (params.mapper == 'minimap2')        { minimap2(combined_ch)   ; mapperout = minimap2.out.fasta  ; mapperbam = minimap2.out.bam}
    else if (params.mapper == 'bwa')        { bwa(combined_ch)        ; mapperout = bwa.out.fasta       ; mapperbam = bwa.out.bam }   
    else                                    { minimap2(combined_ch)   ; mapperout = minimap2.out.fasta  ; mapperbam = minimap2.out.bam}


    bedtools(mapperout)
    plasflow_degen(bedtools.out.fasta)


    // report masked status
    report_ch = plasflow_degen.out.report.view { name, N, W, S, M, K, R, Y, B, D, H, V -> "$name (chromosome): N:$N, W:$W, S:$S, M:$M, K:$K, R:$R, Y:$Y, B:$B, D:$D, H:$H, V:$V" }


    report_write_ch = bedtools.out.report.join(plasflow_degen.out.report)
        .collectFile(seed: 'name,type,N(ATCG),W(AT),S(CG),M(AC),K(TG),R(AG),Y(TC),B(TCG),D(ATG),H(ATC),V(ACG)\n', 
                    storeDir: params.output + "/") {
                    row -> [ "masked_bases_summary.csv", row[0] + ',' + 'genome' + "," + row[1] + ',' + row[2] + "," + row[3] + ',' + row[4] + ',' +
                    row[5] + ',' + row[6] + ',' + row[7] + ',' + row[8] + ',' + row[9] + ',' + row[10] + ',' + row[11] + ',' + '\n' +
                    row[0] + ',' + 'chromosome' + "," + row[12] + ',' + row[13] + "," + row[14] + ',' + row[15] + ',' +
                    row[16] + ',' + row[17] + ',' + row[18] + ',' + row[19] + ',' + row[20] + ',' + row[21] + ',' + row[22] + ',' + '\n'
                    ]
                    }

    emit: 
        plasflow_degen.out.chromosomes
        mapperbam
}

/*
workflow mask_regions_wf {
    take:   
        fastq
        fasta
    main: 

    // join channels
    combined_ch = fasta.join(fastq, by:0)

    // check if everything is alright
    no_match = combined_ch.ifEmpty{ log.info "\033[0;33mCould not match any reads to genomes, please read the help via --help\033[0m" }

    minimap2(combined_ch)
    plasflow(minimap2.out.fasta)

    // report masked status
    report_ch = plasflow.out.report.view { name, all, chr -> "$name: Total masked Bases $all with $chr on the chromosome only." }

    report_write_ch = plasflow.out.report
        .collectFile(seed: 'name,total masked bases,masked bases chromosome\n', 
                    storeDir: params.output + "/") {
                    row -> [ "masked_bases_summary.csv", row[0] + ',' + row[1] + ',' + row[2] + '\n']
                    }

    emit: plasflow.out.chromosomes


}
*/
```
