## Supplementary Code 2 for "Accurate bacterial outbreak tracing with Oxford Nanopore sequencing and reduction of methylation-induced errors": bedtools.nf

```
process bedtools {
        label 'bedtools'
        publishDir "${params.output}/${name}/", mode: 'copy'
    input:
        tuple val(name), path(fasta), path(depths)
  	output:
    	tuple val(name), file("${name}.masked.dm.fasta"), emit: fasta
        tuple val(name), env(BASE_N), env(BASE_W), env(BASE_S), env(BASE_M), env(BASE_K), env(BASE_R), 
                         env(BASE_Y), env(BASE_B), env(BASE_D), env(BASE_H), env(BASE_V), emit: report
  	script:
    """
    # remove lines by depth below params and create bed file
    awk -F'\\t' '\$3<${params.depth}' ${depths} | awk  -F'\\t' '{ printf "%s\\t%s\\t%s\\n", \$1,\$2,\$2 }' > masking.bed

    # mask based on bed file
    bedtools maskfasta -fi ${fasta} -bed masking.bed -fo ${name}.masked.dm.fasta

    # get Masked Bases
    BASE_N=\$(grep -v ">" ${name}.masked.dm.fasta | grep -o "N" | wc -l)
    BASE_W=\$(grep -v ">" ${name}.masked.dm.fasta | grep -o "W" | wc -l)
    BASE_S=\$(grep -v ">" ${name}.masked.dm.fasta | grep -o "S" | wc -l)
    BASE_M=\$(grep -v ">" ${name}.masked.dm.fasta | grep -o "M" | wc -l)
    BASE_K=\$(grep -v ">" ${name}.masked.dm.fasta | grep -o "K" | wc -l)
    BASE_R=\$(grep -v ">" ${name}.masked.dm.fasta | grep -o "R" | wc -l)
    BASE_Y=\$(grep -v ">" ${name}.masked.dm.fasta | grep -o "Y" | wc -l)
    BASE_B=\$(grep -v ">" ${name}.masked.dm.fasta | grep -o "B" | wc -l)
    BASE_D=\$(grep -v ">" ${name}.masked.dm.fasta | grep -o "D" | wc -l)
    BASE_H=\$(grep -v ">" ${name}.masked.dm.fasta | grep -o "H" | wc -l)
    BASE_V=\$(grep -v ">" ${name}.masked.dm.fasta | grep -o "V" | wc -l)
    """
    stub:
    """
    touch ${name}_masked.fasta ${name}.masked.sorted.bam
    BASE_N=5
    BASE_W=7
    BASE_S=1
    BASE_M=3
    BASE_K=0
    BASE_R=0
    BASE_Y=0
    BASE_B=20
    BASE_D=15
    BASE_H=3
    BASE_V=0
    """
}
```
