## Supplementary Code 2 for "Accurate bacterial outbreak tracing with Oxford Nanopore sequencing and reduction of methylation-induced errors": bowtie2.nf

```
process bowtie2 {
        label 'bowtie2'
        publishDir "${params.output}/${name}/", mode: 'copy'
    input:
        tuple val(name), path(fasta), path(reads)
  	output:
    	tuple val(name), file("${name}.masked.fasta"), env(MASKREGIONS), emit: fasta
        tuple val(name), file("${name}.masked.sorted.bam"), emit: bam
  	script:
    if (params.keep_coordinates) {
    """
    bowtie2-build --threads ${task.cpus} ${fasta} ${fasta}.index
    bowtie2 -p ${task.cpus} -f -x ${fasta}.index -U ${reads} | samtools view -bS | samtools sort - -@ ${task.cpus} -o ${name}.minimap.sorted.bam

    # consensus
    samtools consensus -aa --show-ins no --show-del no -@ ${task.cpus} -f fasta -X r10.4_sup ${name}.minimap.sorted.bam -o ${name}.masked.fasta
    rm ${name}.minimap.sorted.bam ${name}.sam

    # get Masked Bases
    MASKREGIONS=\$(grep -v ">" ${name}.masked.fasta | grep -o "N" | wc -l)

    # rebam to masked file for visuals
    minimap2 -t ${task.cpus} -o ${name}.masked.sam -ax map-ont ${name}.masked.fasta ${reads}
    samtools view -bS ${name}.masked.sam | samtools sort - -@ ${task.cpus} -o ${name}.masked.sorted.bam

    rm ${name}.masked.sam
    """
    }
    else {
    """
    bowtie2-build --threads ${task.cpus} ${fasta} ${fasta}.index
    bowtie2 -p ${task.cpus} -f -x ${fasta}.index -U ${reads} | samtools view -bS | samtools sort - -@ ${task.cpus} -o ${name}.minimap.sorted.bam

    # consensus
    samtools consensus -@ ${task.cpus} -f fasta -X r10.4_sup ${name}.minimap.sorted.bam -o ${name}.masked.fasta
    rm ${name}.minimap.sorted.bam ${name}.sam

    # get Masked Bases
    MASKREGIONS=\$(grep -v ">" ${name}.masked.fasta | grep -o "N" | wc -l)

    # rebam to masked file for visuals
    minimap2 -t ${task.cpus} -o ${name}.masked.sam -ax map-ont ${name}.masked.fasta ${reads}
    samtools view -bS ${name}.masked.sam | samtools sort - -@ ${task.cpus} -o ${name}.masked.sorted.bam

    rm ${name}.masked.sam
    """
    }
    stub:
    """
    touch ${name}_masked.fasta
    MASKREGIONS="500"
    """
}

process bowtie2_degen {
        label 'bowtie2'
        publishDir "${params.output}/${name}/", mode: 'copy'
    input:
        tuple val(name), path(fasta), path(reads)
  	output:
    	tuple val(name), path("${name}.masked.fasta"), path("${name}_depth_file.txt"), emit: fasta
        tuple val(name), path("${name}.masked.sorted.bam"), emit: bam
  	script:
    if (params.keep_coordinates) {
    """
    bowtie2-build --threads ${task.cpus} ${fasta} ${fasta}.index
    bowtie2 -p ${task.cpus} -f -x ${fasta}.index -U ${reads} | samtools view -bS | samtools sort - -@ ${task.cpus} -o ${name}.minimap.sorted.bam

    # consensus
    samtools consensus -aa --show-ins no --show-del no \
                        -@ ${task.cpus} \
                        --ambig \
                        -f fasta  \
                        -X r10.4_sup \
                        ${name}.minimap.sorted.bam \
                        -o ${name}.masked.fasta


    # reduce disk footprint                    
    rm ${name}.minimap.sorted.bam ${name}.sam

    # rebam to masked file for visuals
    minimap2 -t ${task.cpus} -o ${name}.masked.sam -ax map-ont ${name}.masked.fasta ${reads}
    samtools view -bS ${name}.masked.sam | samtools sort - -@ ${task.cpus} -o ${name}.masked.sorted.bam

    # get depth per position
    samtools depth -a ${name}.masked.sorted.bam > ${name}_depth_file.txt

    # reduce disk footprint                    
    rm ${name}.masked.sam
    """
    } else {
    """
    bowtie2-build --threads ${task.cpus} ${fasta} ${fasta}.index
    #bowtie2 -p ${task.cpus} -f -x ${fasta}.index -U ${reads} | samtools view -bS | samtools sort - -@ ${task.cpus} -o ${name}.minimap.sorted.bam
    zcat ${reads} > reads_unpacked.fastq
    bowtie2 -p ${task.cpus} -f -x ${fasta}.index -U reads_unpacked.fastq # | samtools view -bS | samtools sort > ${name}.minimap.sorted.bam

    # consensus
    samtools consensus -@ ${task.cpus} \
                        --ambig \
                        -f fasta  \
                        -X r10.4_sup \
                        ${name}.minimap.sorted.bam \
                        -o ${name}.masked.fasta


    # reduce disk footprint                    
    rm ${name}.minimap.sorted.bam ${name}.sam

    # rebam to masked file for visuals
    minimap2 -t ${task.cpus} -o ${name}.masked.sam -ax map-ont ${name}.masked.fasta ${reads}
    samtools view -bS ${name}.masked.sam | samtools sort - -@ ${task.cpus} -o ${name}.masked.sorted.bam

    # get depth per position
    samtools depth -a ${name}.masked.sorted.bam > ${name}_depth_file.txt

    # reduce disk footprint                    
    rm ${name}.masked.sam
    """
    }
    stub:
    """
    touch ${name}_masked.fasta ${name}.masked.sorted.bam
    BASE_N=5
    BASE_W=7
    BASE_S=1
    BASE_M=3
    BASE_K=0
    BASE_R=0
    BASE_Y=0
    BASE_B=20
    BASE_D=15
    BASE_H=3
    BASE_V=0
    """
}
```
