## Supplementary Code 2 for "Accurate bacterial outbreak tracing with Oxford Nanopore sequencing and reduction of methylation-induced errors": count_positions.nf

```
process count_positions {
        label 'seqkit'
        publishDir "${params.output}/${name}/SNP_positions/", mode: 'copy'
    input:
        tuple val(name), path(fasta)
    output:
        tuple val(name), path(fasta), path("*_positions.txt"), optional: true
    script:
        """
        grep -o . <<< "WSMKRYBDHV" | while read BASE; do 
                seqkit locate --ignore-case --only-positive-strand  -r --pattern "\${BASE}" ${fasta} > \${BASE}_positions.txt

                if [ "\$( wc -l <\${BASE}_positions.txt )" -eq 1 ]; then
                    rm -f \${BASE}_positions.txt
                fi
        done
        """
}
```
