## Supplementary Code 2 for "Accurate bacterial outbreak tracing with Oxford Nanopore sequencing and reduction of methylation-induced errors": extract_sequence.nf

```
process extract_sequence {
        label 'samtools'
        publishDir "${params.output}/${name}/SNP_regions/", mode: 'copy'
        errorStrategy 'ignore'
    input:
        tuple val(name), path(fasta), path(positions)
    output:
        tuple val(name), path("*.regions.fasta"), optional: true
    script:
        """
        # WSMKRYBDHV
        grep -o . <<< "WSMKRYBDHV" | while read BASE; do 
            awk -v  s=${params.motif} '{print \$1, \$4, \$5-s, \$6+s}' \${BASE}_positions.txt | tail -n+2 | tr "+" ":" | sed 's/ //'| sed 's/ //' | tr " " "-" > \${BASE}_input_samtools.tsv
            
            # if file empty
            if [ -s \${BASE}_input_samtools.tsv ]; then
                    # The file is not-empty.
                    samtools faidx ${fasta} -c -r \${BASE}_input_samtools.tsv > \${BASE}.regions.fasta
            else
                    # The file is empty.
                    echo "skipping samtools for \${BASE}"
            fi

        done
        """
}
```
