## Supplementary Code 2 for "Accurate bacterial outbreak tracing with Oxford Nanopore sequencing and reduction of methylation-induced errors": get_frequency.nf

```
process get_frequency {
        errorStrategy 'ignore'
        label 'frequency'
        publishDir "${params.output}/${name}/pileup/", mode: 'copy'
    input:
        tuple val(name), path(fasta), path(positions), path(bam) 

  	output:
    	tuple val(name), file("${name}_Frequency.tsv"), path("${name}_pileup.tsv")
  	script:
    """
    # ACTIVATE HISTORY
        set -euxo pipefail

    # remove headers and create one correct bed position file for samtools (name \t position)
    tail -q -n+2 ${positions}  | awk -v OFS='\\t' '{print \$1,\$5}' > clean.bed

    # pileup for all positions
    samtools index ${bam}
    samtools mpileup ${bam} -f ${fasta} --positions clean.bed --output ${name}_pileup.tsv

    # count stuff per degenerate base
    ## Y
    YT_COUNT=\$(awk -F'\\t' '{if(\$3=="Y")print \$5}' ${name}_pileup.tsv | sed 's/[^T]//g' | awk '{ print length }')
    YC_COUNT=\$(awk -F'\\t' '{if(\$3=="Y")print \$5}' ${name}_pileup.tsv | sed 's/[^C]//g' | awk '{ print length }')   
    Yt_COUNT=\$(awk -F'\\t' '{if(\$3=="Y")print \$5}' ${name}_pileup.tsv | sed 's/[^t]//g' | awk '{ print length }')
    Yc_COUNT=\$(awk -F'\\t' '{if(\$3=="Y")print \$5}' ${name}_pileup.tsv | sed 's/[^c]//g' | awk '{ print length }')    

    Y_TOTAL=\$(awk -F'\\t' '{if(\$3=="Y")print \$5}' ${name}_pileup.tsv | sed 's/[^CT]//g' | awk '{ print length }')
    y_TOTAL=\$(awk -F'\\t' '{if(\$3=="Y")print \$5}' ${name}_pileup.tsv | sed 's/[^ct]//g' | awk '{ print length }')    

    paste -d'\\t' <(echo "\$Y_TOTAL") <(echo "\$YT_COUNT") | awk -v OFS='\\t' '{print \$2 / \$1}'| sed 's/^/T\\t/' | sed 's/^/Y\\t/' | sed 's/^/forward\\t/' >> ${name}_Frequency.tsv
    paste -d'\\t' <(echo "\$Y_TOTAL") <(echo "\$YC_COUNT") | awk -v OFS='\\t' '{print \$2 / \$1}'| sed 's/^/C\\t/' | sed 's/^/Y\\t/' | sed 's/^/forward\\t/' >> ${name}_Frequency.tsv
    paste -d'\\t' <(echo "\$y_TOTAL") <(echo "\$Yt_COUNT") | awk -v OFS='\\t' '{print \$2 / \$1}'| sed 's/^/T\\t/' | sed 's/^/Y\\t/' | sed 's/^/reverse\\t/' >> ${name}_Frequency.tsv
    paste -d'\\t' <(echo "\$y_TOTAL") <(echo "\$Yc_COUNT") | awk -v OFS='\\t' '{print \$2 / \$1}'| sed 's/^/C\\t/' | sed 's/^/Y\\t/' | sed 's/^/reverse\\t/' >> ${name}_Frequency.tsv

    # unsetting, might need to do this earlier in batches depending on the RAM
    unset YT_COUNT YC_COUNT Yt_COUNT Yc_COUNT Y_TOTAL y_TOTAL

    ## R
    RA_COUNT=\$(awk -F'\\t' '{if(\$3=="R")print \$5}' ${name}_pileup.tsv | sed 's/[^A]//g' | awk '{ print length }')
    RG_COUNT=\$(awk -F'\\t' '{if(\$3=="R")print \$5}' ${name}_pileup.tsv | sed 's/[^G]//g' | awk '{ print length }')    
    Ra_COUNT=\$(awk -F'\\t' '{if(\$3=="R")print \$5}' ${name}_pileup.tsv | sed 's/[^a]//g' | awk '{ print length }')
    Rg_COUNT=\$(awk -F'\\t' '{if(\$3=="R")print \$5}' ${name}_pileup.tsv | sed 's/[^g]//g' | awk '{ print length }')  

    R_TOTAL=\$(awk -F'\\t' '{if(\$3=="R")print \$5}' ${name}_pileup.tsv | sed 's/[^AG]//g' | awk '{ print length }')
    r_TOTAL=\$(awk -F'\\t' '{if(\$3=="R")print \$5}' ${name}_pileup.tsv | sed 's/[^ag]//g' | awk '{ print length }')    
    
    paste -d'\\t' <(echo "\$R_TOTAL") <(echo "\$RA_COUNT") | awk -v OFS='\\t' '{print \$2 / \$1}'| sed 's/^/A\\t/' | sed 's/^/R\\t/' | sed 's/^/forward\\t/' >> ${name}_Frequency.tsv
    paste -d'\\t' <(echo "\$R_TOTAL") <(echo "\$RG_COUNT") | awk -v OFS='\\t' '{print \$2 / \$1}'| sed 's/^/G\\t/' | sed 's/^/R\\t/' | sed 's/^/forward\\t/' >> ${name}_Frequency.tsv
    paste -d'\\t' <(echo "\$r_TOTAL") <(echo "\$Ra_COUNT") | awk -v OFS='\\t' '{print \$2 / \$1}'| sed 's/^/A\\t/' | sed 's/^/R\\t/' | sed 's/^/reverse\\t/' >> ${name}_Frequency.tsv
    paste -d'\\t' <(echo "\$r_TOTAL") <(echo "\$Rg_COUNT") | awk -v OFS='\\t' '{print \$2 / \$1}'| sed 's/^/G\\t/' | sed 's/^/R\\t/' | sed 's/^/reverse\\t/' >> ${name}_Frequency.tsv

    unset RA_COUNT RG_COUNT Ra_COUNT Rg_COUNT R_TOTAL r_TOTAL

    ## W
    WA_COUNT=\$(awk -F'\\t' '{if(\$3=="W")print \$5}' ${name}_pileup.tsv | sed 's/[^A]//g' | awk '{ print length }')
    WT_COUNT=\$(awk -F'\\t' '{if(\$3=="W")print \$5}' ${name}_pileup.tsv | sed 's/[^T]//g' | awk '{ print length }')    
    Wa_COUNT=\$(awk -F'\\t' '{if(\$3=="W")print \$5}' ${name}_pileup.tsv | sed 's/[^a]//g' | awk '{ print length }')
    Wt_COUNT=\$(awk -F'\\t' '{if(\$3=="W")print \$5}' ${name}_pileup.tsv | sed 's/[^t]//g' | awk '{ print length }') 

    W_TOTAL=\$(awk -F'\\t' '{if(\$3=="W")print \$5}' ${name}_pileup.tsv | sed 's/[^AT]//g' | awk '{ print length }')
    w_TOTAL=\$(awk -F'\\t' '{if(\$3=="W")print \$5}' ${name}_pileup.tsv | sed 's/[^at]//g' | awk '{ print length }')
    
    paste -d'\\t' <(echo "\$W_TOTAL") <(echo "\$WA_COUNT") | awk -v OFS='\\t' '{print \$2 / \$1}'| sed 's/^/A\\t/' | sed 's/^/W\\t/' | sed 's/^/forward\\t/' >> ${name}_Frequency.tsv
    paste -d'\\t' <(echo "\$W_TOTAL") <(echo "\$WT_COUNT") | awk -v OFS='\\t' '{print \$2 / \$1}'| sed 's/^/T\\t/' | sed 's/^/W\\t/' | sed 's/^/forward\\t/' >> ${name}_Frequency.tsv
    paste -d'\\t' <(echo "\$w_TOTAL") <(echo "\$Wa_COUNT") | awk -v OFS='\\t' '{print \$2 / \$1}'| sed 's/^/A\\t/' | sed 's/^/W\\t/' | sed 's/^/reverse\\t/' >> ${name}_Frequency.tsv
    paste -d'\\t' <(echo "\$w_TOTAL") <(echo "\$Wt_COUNT") | awk -v OFS='\\t' '{print \$2 / \$1}'| sed 's/^/T\\t/' | sed 's/^/W\\t/' | sed 's/^/reverse\\t/' >> ${name}_Frequency.tsv    

    unset WA_COUNT WT_COUNT Wa_COUNT Wt_COUNT W_TOTAL w_TOTAL

    ## S
    SG_COUNT=\$(awk -F'\\t' '{if(\$3=="S")print \$5}' ${name}_pileup.tsv | sed 's/[^G]//g' | awk '{ print length }')
    SC_COUNT=\$(awk -F'\\t' '{if(\$3=="S")print \$5}' ${name}_pileup.tsv | sed 's/[^C]//g' | awk '{ print length }')    
    Sg_COUNT=\$(awk -F'\\t' '{if(\$3=="S")print \$5}' ${name}_pileup.tsv | sed 's/[^g]//g' | awk '{ print length }')
    Sc_COUNT=\$(awk -F'\\t' '{if(\$3=="S")print \$5}' ${name}_pileup.tsv | sed 's/[^c]//g' | awk '{ print length }')  

    S_TOTAL=\$(awk -F'\\t' '{if(\$3=="S")print \$5}' ${name}_pileup.tsv | sed 's/[^CG]//g' | awk '{ print length }')
    s_TOTAL=\$(awk -F'\\t' '{if(\$3=="S")print \$5}' ${name}_pileup.tsv | sed 's/[^cg]//g' | awk '{ print length }')    
    
    paste -d'\\t' <(echo "\$S_TOTAL") <(echo "\$SG_COUNT") | awk -v OFS='\\t' '{print \$2 / \$1}'| sed 's/^/G\\t/' | sed 's/^/S\\t/' | sed 's/^/forward\\t/' >> ${name}_Frequency.tsv
    paste -d'\\t' <(echo "\$S_TOTAL") <(echo "\$SC_COUNT") | awk -v OFS='\\t' '{print \$2 / \$1}'| sed 's/^/C\\t/' | sed 's/^/S\\t/' | sed 's/^/forward\\t/' >> ${name}_Frequency.tsv
    paste -d'\\t' <(echo "\$s_TOTAL") <(echo "\$Sg_COUNT") | awk -v OFS='\\t' '{print \$2 / \$1}'| sed 's/^/G\\t/' | sed 's/^/S\\t/' | sed 's/^/reverse\\t/' >> ${name}_Frequency.tsv
    paste -d'\\t' <(echo "\$s_TOTAL") <(echo "\$Sc_COUNT") | awk -v OFS='\\t' '{print \$2 / \$1}'| sed 's/^/C\\t/' | sed 's/^/S\\t/' | sed 's/^/reverse\\t/' >> ${name}_Frequency.tsv    

    unset SG_COUNT SC_COUNT Sg_COUNT Sc_COUNT S_TOTAL s_TOTAL

    ## M
    MA_COUNT=\$(awk -F'\\t' '{if(\$3=="M")print \$5}' ${name}_pileup.tsv | sed 's/[^A]//g' | awk '{ print length }')
    MC_COUNT=\$(awk -F'\\t' '{if(\$3=="M")print \$5}' ${name}_pileup.tsv | sed 's/[^C]//g' | awk '{ print length }')    
    Ma_COUNT=\$(awk -F'\\t' '{if(\$3=="M")print \$5}' ${name}_pileup.tsv | sed 's/[^a]//g' | awk '{ print length }')
    Mc_COUNT=\$(awk -F'\\t' '{if(\$3=="M")print \$5}' ${name}_pileup.tsv | sed 's/[^c]//g' | awk '{ print length }')  

    M_TOTAL=\$(awk -F'\\t' '{if(\$3=="M")print \$5}' ${name}_pileup.tsv | sed 's/[^AC]//g' | awk '{ print length }')
    m_TOTAL=\$(awk -F'\\t' '{if(\$3=="M")print \$5}' ${name}_pileup.tsv | sed 's/[^ac]//g' | awk '{ print length }')    
    
    paste -d'\\t' <(echo "\$M_TOTAL") <(echo "\$MA_COUNT") | awk -v OFS='\t' '{print \$2 / \$1}'| sed 's/^/A\\t/' | sed 's/^/M\\t/' | sed 's/^/forward\\t/' >> ${name}_Frequency.tsv
    paste -d'\\t' <(echo "\$M_TOTAL") <(echo "\$MC_COUNT") | awk -v OFS='\t' '{print \$2 / \$1}'| sed 's/^/C\\t/' | sed 's/^/M\\t/' | sed 's/^/forward\\t/' >> ${name}_Frequency.tsv
    paste -d'\\t' <(echo "\$m_TOTAL") <(echo "\$Ma_COUNT") | awk -v OFS='\t' '{print \$2 / \$1}'| sed 's/^/A\\t/' | sed 's/^/M\\t/' | sed 's/^/reverse\\t/' >> ${name}_Frequency.tsv
    paste -d'\\t' <(echo "\$m_TOTAL") <(echo "\$Mc_COUNT") | awk -v OFS='\t' '{print \$2 / \$1}'| sed 's/^/C\\t/' | sed 's/^/M\\t/' | sed 's/^/reverse\\t/' >> ${name}_Frequency.tsv

    unset MA_COUNT MC_COUNT Ma_COUNT Mc_COUNT M_TOTAL m_TOTAL

    ## K
    KG_COUNT=\$(awk -F'\\t' '{if(\$3=="K")print \$5}' ${name}_pileup.tsv | sed 's/[^G]//g' | awk '{ print length }')
    KT_COUNT=\$(awk -F'\\t' '{if(\$3=="K")print \$5}' ${name}_pileup.tsv | sed 's/[^T]//g' | awk '{ print length }')
    Kg_COUNT=\$(awk -F'\\t' '{if(\$3=="K")print \$5}' ${name}_pileup.tsv | sed 's/[^g]//g' | awk '{ print length }')
    Kt_COUNT=\$(awk -F'\\t' '{if(\$3=="K")print \$5}' ${name}_pileup.tsv | sed 's/[^t]//g' | awk '{ print length }')  

    K_TOTAL=\$(awk -F'\\t' '{if(\$3=="K")print \$5}' ${name}_pileup.tsv | sed 's/[^TG]//g' | awk '{ print length }')
    k_TOTAL=\$(awk -F'\\t' '{if(\$3=="K")print \$5}' ${name}_pileup.tsv | sed 's/[^tg]//g' | awk '{ print length }')    
    
    paste -d'\\t' <(echo "\$K_TOTAL") <(echo "\$KG_COUNT") | awk -v OFS='\t' '{print \$2 / \$1}'| sed 's/^/G\\t/' | sed 's/^/K\\t/' | sed 's/^/forward\\t/' >> ${name}_Frequency.tsv
    paste -d'\\t' <(echo "\$K_TOTAL") <(echo "\$KT_COUNT") | awk -v OFS='\t' '{print \$2 / \$1}'| sed 's/^/T\\t/' | sed 's/^/K\\t/' | sed 's/^/forward\\t/' >> ${name}_Frequency.tsv
    paste -d'\\t' <(echo "\$k_TOTAL") <(echo "\$Kg_COUNT") | awk -v OFS='\t' '{print \$2 / \$1}'| sed 's/^/G\\t/' | sed 's/^/K\\t/' | sed 's/^/reverse\\t/' >> ${name}_Frequency.tsv
    paste -d'\\t' <(echo "\$k_TOTAL") <(echo "\$Kt_COUNT") | awk -v OFS='\t' '{print \$2 / \$1}'| sed 's/^/T\\t/' | sed 's/^/K\\t/' | sed 's/^/reverse\\t/' >> ${name}_Frequency.tsv    

    unset KG_COUNT KT_COUNT Kg_COUNT Kt_COUNT K_TOTAL k_TOTAL

    """
}
```
