## Supplementary Code 2 for "Accurate bacterial outbreak tracing with Oxford Nanopore sequencing and reduction of methylation-induced errors": plasflow.nf

```
process plasflow {
        label 'plasflow'
        publishDir "${params.output}/${name}/", mode: 'copy'
    input:
        tuple val(name), path(fasta), val(allmasked) 
    output:
        tuple val(name), path("${name}.masked_chromosomes.fasta"), emit: chromosomes  optional true
        tuple val(name), path("${name}.masked_plasmids_plasmids.fasta"), emit: plasmids  optional true
        tuple val(name), path("${name}.masked_unclassified_unclassified.fasta"), emit: unclassified  optional true
        tuple val(name), val(allmasked), env(MASKREGIONS), emit: report
    script:
        """
        PlasFlow.py --input ${fasta} --output ${name}.masked --threshold 0.7

        # remove empty file
        find . -name "*.fasta" -type f -size 0 -print0 | xargs -0 echo rm

        # get Masked Bases
        MASKREGIONS=\$(grep -v ">" ${name}.masked_chromosomes.fasta | grep -o "N" | wc -l)
        """
        stub:
        """
        touch  ${name}.masked_chromosomes.fasta ${name}.masked_plasmids.fasta ${name}.masked_unclassified.fasta
        """
}

process plasflow_degen {
        label 'plasflow'
        publishDir "${params.output}/${name}/", mode: 'copy'
    input:
        tuple val(name), path(fasta)
    output:
        tuple val(name), path("${name}.masked_chromosomes.fasta"), emit: chromosomes  optional true
        tuple val(name), path("${name}.masked_plasmids.fasta"), emit: plasmids  optional true
        tuple val(name), path("${name}.masked_unclassified.fasta"), emit: unclassified  optional true
        tuple val(name), env(BASE_N), env(BASE_W), env(BASE_S), env(BASE_M), env(BASE_K), env(BASE_R), 
                         env(BASE_Y), env(BASE_B), env(BASE_D), env(BASE_H), env(BASE_V), emit: report
    script:
        """
        PlasFlow.py --input ${fasta} --output ${name}.masked --threshold 0.7

        # remove empty file
        find . -name "*.fasta" -type f -size 0 -print0 | xargs -0 echo rm

        # get Masked Bases
        BASE_N=\$(grep -v ">" ${name}.masked_chromosomes.fasta | grep -o "N" | wc -l)
        BASE_W=\$(grep -v ">" ${name}.masked_chromosomes.fasta | grep -o "W" | wc -l)
        BASE_S=\$(grep -v ">" ${name}.masked_chromosomes.fasta | grep -o "S" | wc -l)
        BASE_M=\$(grep -v ">" ${name}.masked_chromosomes.fasta | grep -o "M" | wc -l)
        BASE_K=\$(grep -v ">" ${name}.masked_chromosomes.fasta | grep -o "K" | wc -l)
        BASE_R=\$(grep -v ">" ${name}.masked_chromosomes.fasta | grep -o "R" | wc -l)
        BASE_Y=\$(grep -v ">" ${name}.masked_chromosomes.fasta | grep -o "Y" | wc -l)
        BASE_B=\$(grep -v ">" ${name}.masked_chromosomes.fasta | grep -o "B" | wc -l)
        BASE_D=\$(grep -v ">" ${name}.masked_chromosomes.fasta | grep -o "D" | wc -l)
        BASE_H=\$(grep -v ">" ${name}.masked_chromosomes.fasta | grep -o "H" | wc -l)
        BASE_V=\$(grep -v ">" ${name}.masked_chromosomes.fasta | grep -o "V" | wc -l)

        """
        stub:
        """
        touch  ${name}_chromosomes.fasta ${name}_plasmids.fasta ${name}_unclassified.fasta
        BASE_N=5
        BASE_W=7
        BASE_S=1
        BASE_M=3
        BASE_K=0
        BASE_R=0
        BASE_Y=0
        BASE_B=20
        BASE_D=15
        BASE_H=3
        BASE_V=0
        """
}
```
