## Supplementary Code 2 for "Accurate bacterial outbreak tracing with Oxford Nanopore sequencing and reduction of methylation-induced errors": plot_frequency.nf

```
process plot_frequency {
        label 'ggplot2'
        publishDir "${params.output}", mode: 'copy'
        //errorStrategy 'ignore'
    input:
        path(frequency)
    output:
        path("frequency.svg"), optional: true

    script:
        """
        cat ${frequency} | grep -v "nan" | grep -v "0\$" > input.tsv
        violin_chart.R
        """

        stub:
        """
        touch frequency.svg
        """
}
```
