## Supplementary Code 2 for "Accurate bacterial outbreak tracing with Oxford Nanopore sequencing and reduction of methylation-induced errors": plot_motifs.nf

```
process plot_motifs {
        label 'ggplot2'
        publishDir "${params.output}/motif_logos", mode: 'copy'
        //errorStrategy 'ignore'
    input:
        tuple val(name), path(fasta)
    output:
        tuple val(name), path("${name}.svg"), optional: true

    script:
        """
        find . -name "?.regions.fasta" -exec awk -v x=11 'NR==x{exit 1}' {} \\; -exec rm -f {} \\;
        FILES=\$(ls | grep ".regions.fasta" | head -1)

        if [ ! -z "\${FILES}" ]; then
    
            create_logo.R ${name}
            mv logo.svg ${name}.svg    
        
        fi

        """

        stub:
        """
        touch ${name}.svg
        """
}
```
