## Supplementary figures and images for "Accurate bacterial outbreak tracing with Oxford Nanopore sequencing and reduction of methylation-induced errors"

### mobile_logo.png

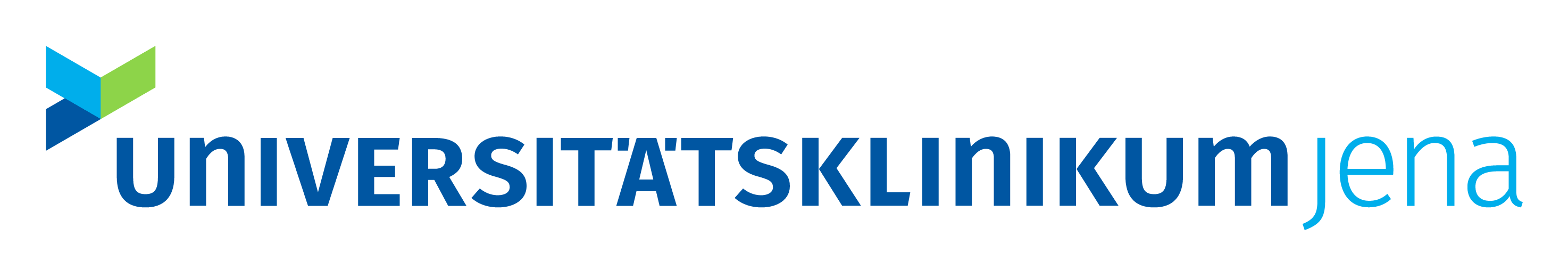

### MPOA_flowchart.png

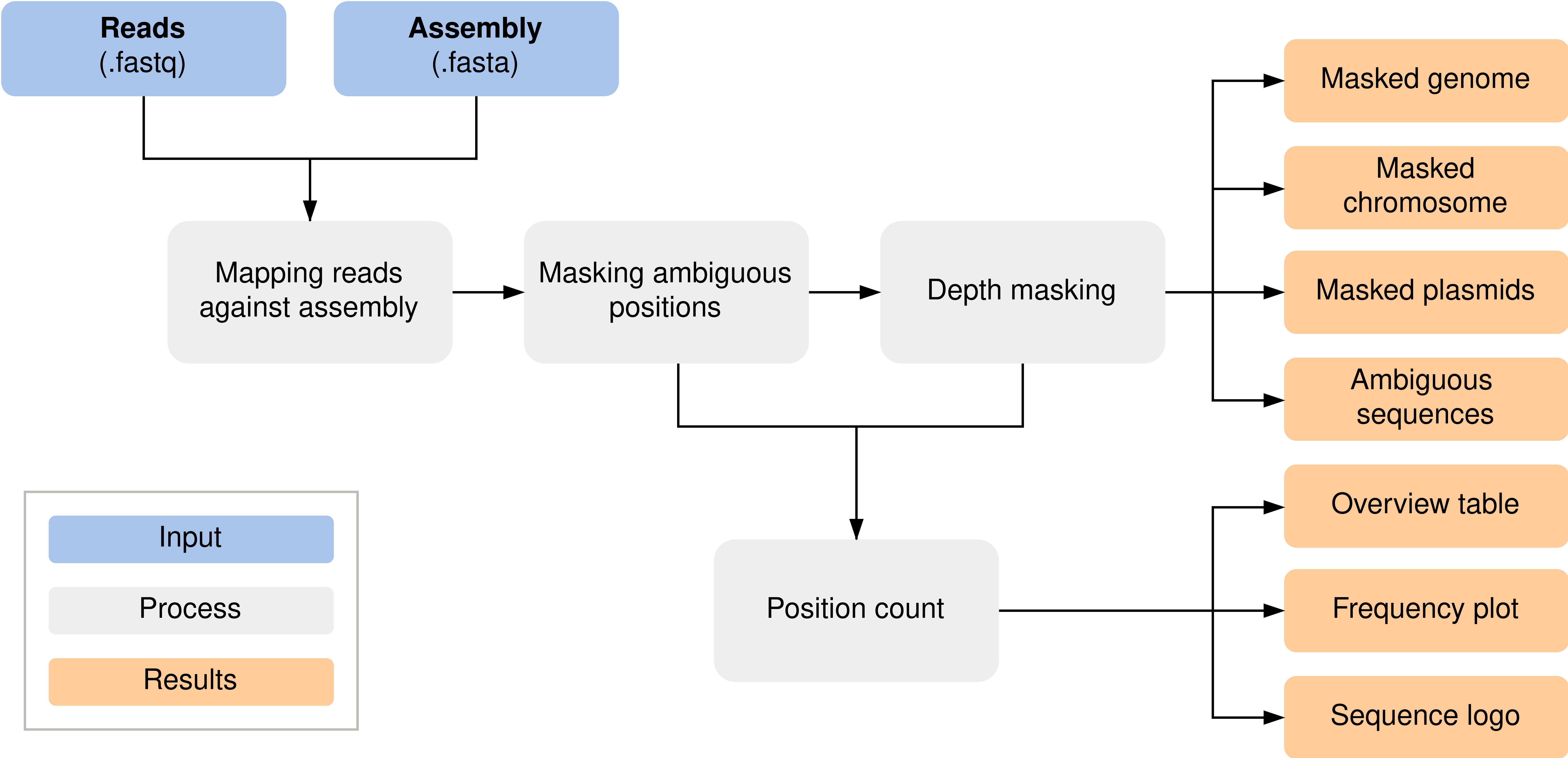
